# Common cardiac medications potently inhibit ACE2 binding to the SARS-CoV-2 Spike, and block virus penetration into human lung cells

**DOI:** 10.1101/2021.06.02.446343

**Authors:** Hung Caohuy, Ofer Eidelman, Tinghua Chen, Qingfeng Yang, Alakesh Bera, Nathan Walton, Harvey B. Pollard

**Author notes:** Communications: Harvey B. Pollard, M.D., Ph.D., Department of Anatomy, Physiology and Genetics Uniformed Services University School of Medicine, Uniformed Services University of the Health Sciences, Bethesda, MD 20814. =co-first authors: these investigators contributed equally to this work (Hung Caohuy, Ofer Eidelman).

## Abstract

To initiate SARS-CoV-2 infection, the Receptor Binding Domain (RBD) on the viral spike protein must first bind to the host receptor ACE2 protein on pulmonary and other ACE2-expressing cells. We hypothesized that cardiac glycoside drugs might block the binding reaction between ACE2 and the Spike (S) protein, and thus block viral penetration into target cells. To test this hypothesis we developed a biochemical assay for ACE2:Spike binding, and tested cardiac glycosides as inhibitors of binding. Here we report that ouabain, digitoxin, and digoxin are high-affinity competitive inhibitors of ACE2 binding to the Wuhan S1 and the European [E614G] S1 proteins. These drugs also inhibit ACE2 binding to the Wuhan RBD, as well as to RBD proteins containing the S. Africa [E484K], Mink [Y453F] and UK [N501Y] mutations. As hypothesized, we also found that ouabain and digitoxin blocked penetration by SARS-CoV-2 Spike-pseudotyped virus into human lung cells. These data indicate that cardiac glycosides may block viral penetration into the target cell by first inhibiting ACE2:Spike binding. Clinical concentrations of ouabain and digitoxin are relatively safe for short term use for subjects with normal hearts. It has therefore not escaped our attention that these common cardiac medications could be deployed worldwide as inexpensive repurposed drugs for anti-COVID-19 therapy.

## Introduction

COVID-19 is a pandemic pneumonia-like disease, caused by the SARS-CoV-2 virus, that was first detected late in 2019 in Wuhan, China ^1–3^. The virus initially gains access to type II pneumocytes in the lung when the Receptor Binding Domain (RBD) on the viral Spike (S) protein binds to the extracellular carboxypeptidase domain of the ACE2 receptor protein on the cell surface ^4–6^. The native spike structure is composed of three identical protomers. Each protomer is divided into two domains: the distal N-terminal S1 spike “tip”, containing the RBD, and the residual C-terminal S2, transmembrane fusion protein. In the intact Spike, the binding process appears to depend on the RBD of at least one protomer transitioning from a down “closed” state to an upper “open” state ^7,8^. In the down “closed” state the RBD has been shown by Cryo-EM to be partially hidden by glycosylated spike amino acids ^9^. When activated by ACE2, the RBD domain emerges towards the spike tip, from the hidden state, into the “open” state. Thus from the perspective of ACE2, there may be at least two accessible conformations of the RBD: either a lower, mostly hidden structure, or a raised, much less hidden structure ^7^. However, how ACE2 induces the transition of the RBD from a closed state to the open state is presently not yet understood.

Following binding of ACE2 to the Spike RBD, the Spike protein is cleaved by Furin, a host protease, into the distal S1 segment, and the residual S2 segment ^10,11^. Furin cleavage reveals the C-terminal *CendR* domain, to which the host protein neuropilin 1 binds and promotes cell entry ^10^. The residual S2 domain is further sequentially optimized for membrane fusion by the host proteases TMPRSS2 and endosomal CTSL ^6,9^. However, since the Wuhan viral strain was first sequenced, the virus has mutated into more infectious and possibly more lethal strains. Examples of such mutations in the spike include the European mutation at residue 614, [D614G] ^12,13^. This mutation appears to increase spike density and flexibility ^14,15^. The mutation also creates an additional cleavage site for serine elastase 2 (neutrophil elastase) that promotes further exposure and priming of S2 ^6^. Additional mutations have been detected in the RBD, including [N501Y], originating in the United Kingdom ^16^; [Y453F] the Denmark/Mink mutation ^17^; and [E484K], a mutation originating in South Africa ^18^. The [E484K] mutation contributes to recent COVID-19 outbreaks in India in the B.1.116 lineage, and in Brazil as variants P.1 and P.2 within the B.1.1.33 lineage. Since infection depends on successful viral entry into the cell, this complex entry mechanism has therefore become a promising target for anti-viral SARS-CoV-2 drug discovery.

In an effort to jumpstart COVID-19 drug discovery, unbiased *in vitro* and *in silico* screens of FDA-approved and other drugs have been performed to find possible high affinity blockers of SARS-CoV-2 infectivity ^19–21^. In some of these screens the common cardiac medications digitoxin, digoxin and ouabain, have emerged as top contenders. Experimentally, time-of addition experiments with infection by native SARS-CoV-2 of green monkey kidney Vero cells have been interpreted to suggest that ouabain blocks virus penetration, while digoxin blocks infectivity at an unknown intracellular site ^22^. Experiments with digoxin ^23^ and digitoxin ^24^ have also been interpreted as blocking entry of native MERS-CoV into target cells. Consistently, *in silico* docking sites for digitoxin have been identified in the RBD ^25^. However, other *in silico* studies have shown that cardiac glycoside drugs might also interact with the active site of the viral main protease, Mpro ^26^. Thus it remains to be determined where in the infectivity process these cardiac glycoside drugs act, since in most studies the experimental endpoints have been viral progeny counts.

Based on the foregoing we hypothesized that cardiac glycoside drugs such as ouabain, digitoxin and digoxin might block the binding reaction between ACE2 and the viral Spike (S) protein, and thus block viral penetration into target cells (**Supplemental Figure S1**). To test this hypothesis we first developed a biochemical method to measure the kinetics of ACE2 binding to immobilized variants of the Spike S1 and RBD domains. We found that binding occurred by a positively cooperative mechanism. Next we tested the cardiac glycoside drugs for their ability to block the ACE2:Spike binding reaction. These drugs were all found to be competitive inhibitors. Lastly we tested whether cardiac glycoside drugs could block viral entry into human lung cells with SARS-CoV-2-spike-pseudotyped virus. The most potent blocker was ouabain, followed by digitoxin. The least potent was digoxin. These cardiac glycoside drugs are inexpensive, widely available, and clinically safe for subjects with normal hearts ^27–29^. It is therefore possible that these drugs could be repurposed for COVID-19 prevention and therapy.

## Results

### ACE2 binds with positive cooperativity to Receptor Binding Domain proteins

The substrate-binding plots in **Figure 1a** show that both the original Wuhan RBD protein, and the larger original Wuhan [D614] S1 domain protein, appear to have very similar binding isotherms at 37°C. However, the mutant European [D614G] S1 differs quantitatively from the original Wuhan S1 and Wuhan RBD. *Furthermore, inspection of the substrate-binding plots show that a lag in binding occurs at low concentrations of ACE2. Thus, the binding isotherms appear to show a distinct sigmoid appearance in the low concentration range that is independent of mutation.* To further test this possibility, we constructed Eadie-Hofstee (EH) plots for each of the domains. In the case of the original Wuhan [D614] S1 protein **Figure 1b** shows a “concave-down” structure at the lower values of ACE2 binding. Such behavior is characteristic of positive cooperativity, including positive cooperativity by monomeric proteins or enzymes ^30^. **Figure 1c** and **Figure 1d** show the same descriptive positive cooperativity behavior in Eadie-Hofstee plots for both the mutant [D614G] S1 protein and the Wuhan RBD protein, respectively. Hill plots of these data also show the modest but significant elevations in the Hill coefficient, (n_H_) which is typical of monomeric systems ^30^ (**Supplementary Figure S5**). It is apparent that since both Spike S1 proteins contain the same upstream RBD domain, the occurrence of either the amino acid aspartic acid (D) or glycine (G) at residue 614 has the ability to modify ACE2 affinity for the common RBD. *The data further indicate that the positive cooperativity property may be intrinsic to the ACE2 interaction with the RBD*. The values of K_D_ for each variant are included in **Table 1**.

**Figure 1.**
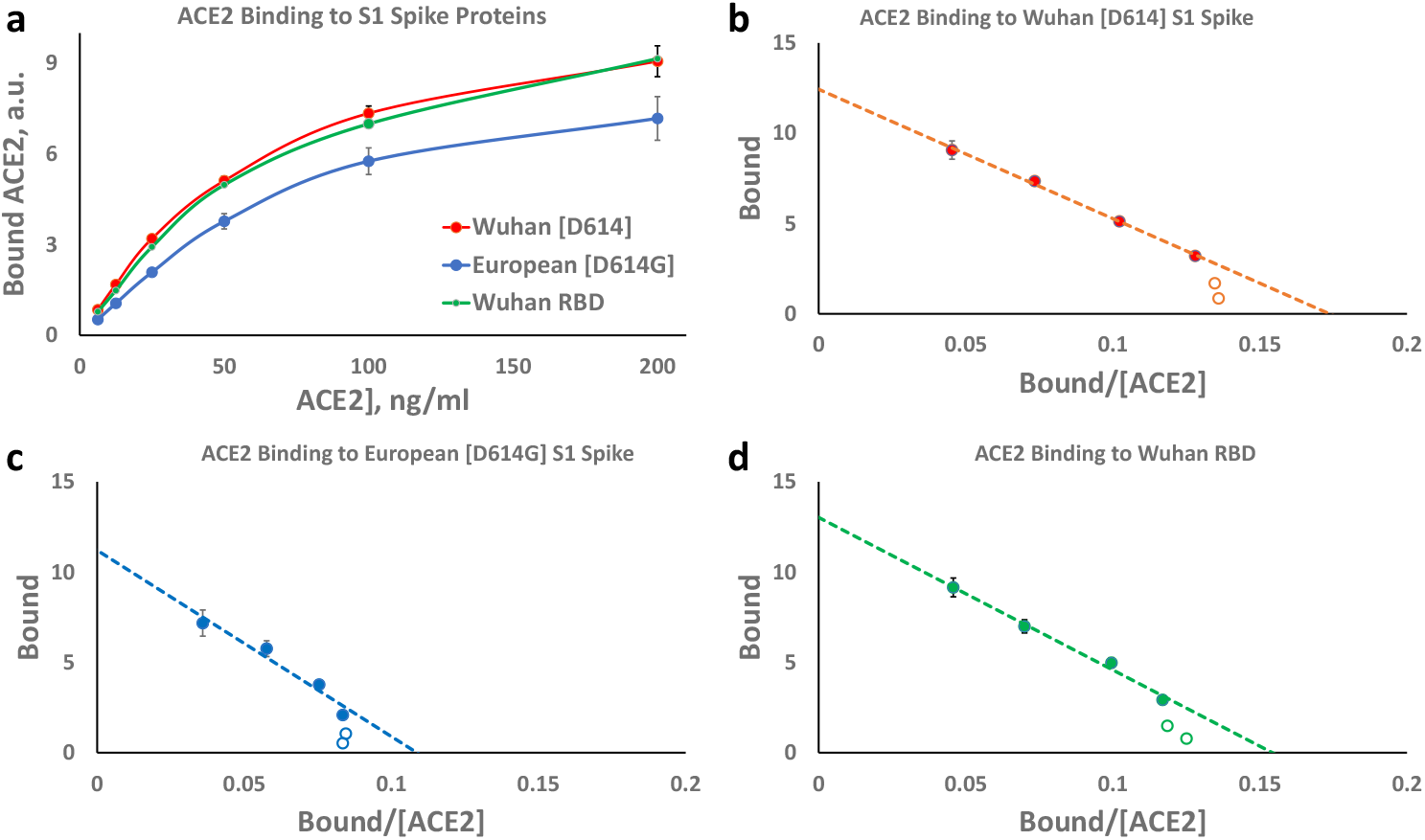
Binding of ACE2 to the SARS-CoV-2 Spike variants. (**a**) Substrate-Binding plot for the original Wuhan S1 (red), the European mutant [D614G] S1 (blue), and the Wuhan RBD (green) proteins. (**b**) Eadie-Hoffstee plot for Wuhan [D614] S1 spike. Note concave-down structure at low binding levels. (**c**) Eadie-Hoffstee plot for European [D614G] S1 spike protein. (**d**) Eadie-Hoffstee plot for Wuhan RBD from the S1 spike. Individual points are the average of quadruplicate technical replicas from 6 independent experiments. Each point is the average ± SE for 5-6 independent experiments.

**Table 1.**
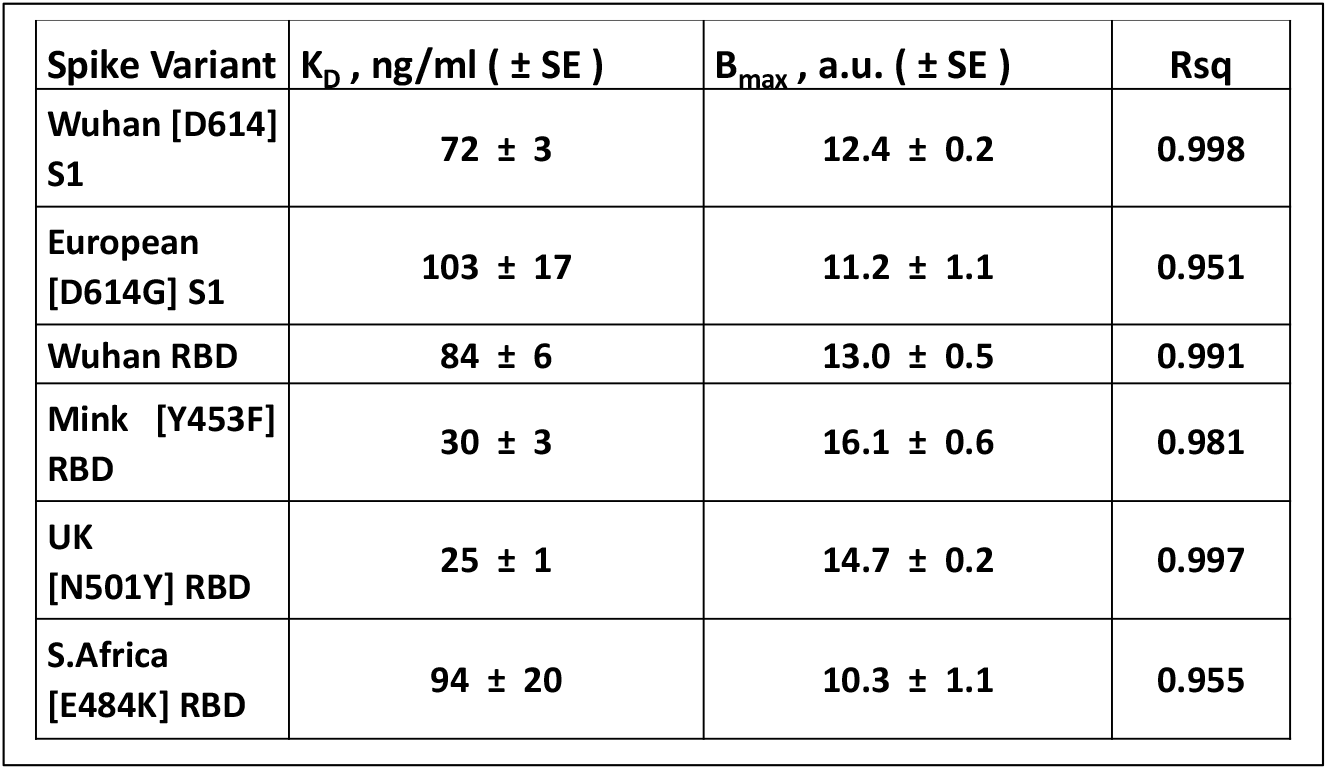
Binding constants K_D_ for ACE2 binding to spike variants.

We also tested ACE2 binding properties to RBD proteins containing three other disease-associated mutations, and compared them to the original Wuhan RBD. **Figure 2a** shows the substrate-binding isotherms for the Mink [Y453F] RBD and the UK [N501Y] RBD proteins, and compares them with the Wuhan S1, and the Wuhan RBD proteins. **Figure 2b** shows the Eadie-Hoffstee plots for the Wuhan S1 and the Wuhan RBD. The Wuhan RBD domain is contained within the Wuhan S1 protein, and as expected the data are virtually superimposable. Both curves are concave down at low ACE2 binding levels, and thus are consistent with the interpretation of positive cooperativity. **Figures 2c** and **2d** show the same type of data for the Mink [Y453F] RBD protein and the UK [N501Y] RBD protein, respectively. The Eadie-Hoffstee plots are concave down at low values of binding, while the slopes in the linear parts of the plots are substantially *reduced* compared to the Wuhan S1 proteins. As summarized in **Table 1**, K_D_ values for the Mink and UK variants are significantly reduced from 72 to 30 and 25 ng/ml ACE2, respectively*. Thus mutations in the RBD in the region interacting with ACE2 can affect the K*_*D*_ *for the interaction.*

**Figure 2.**
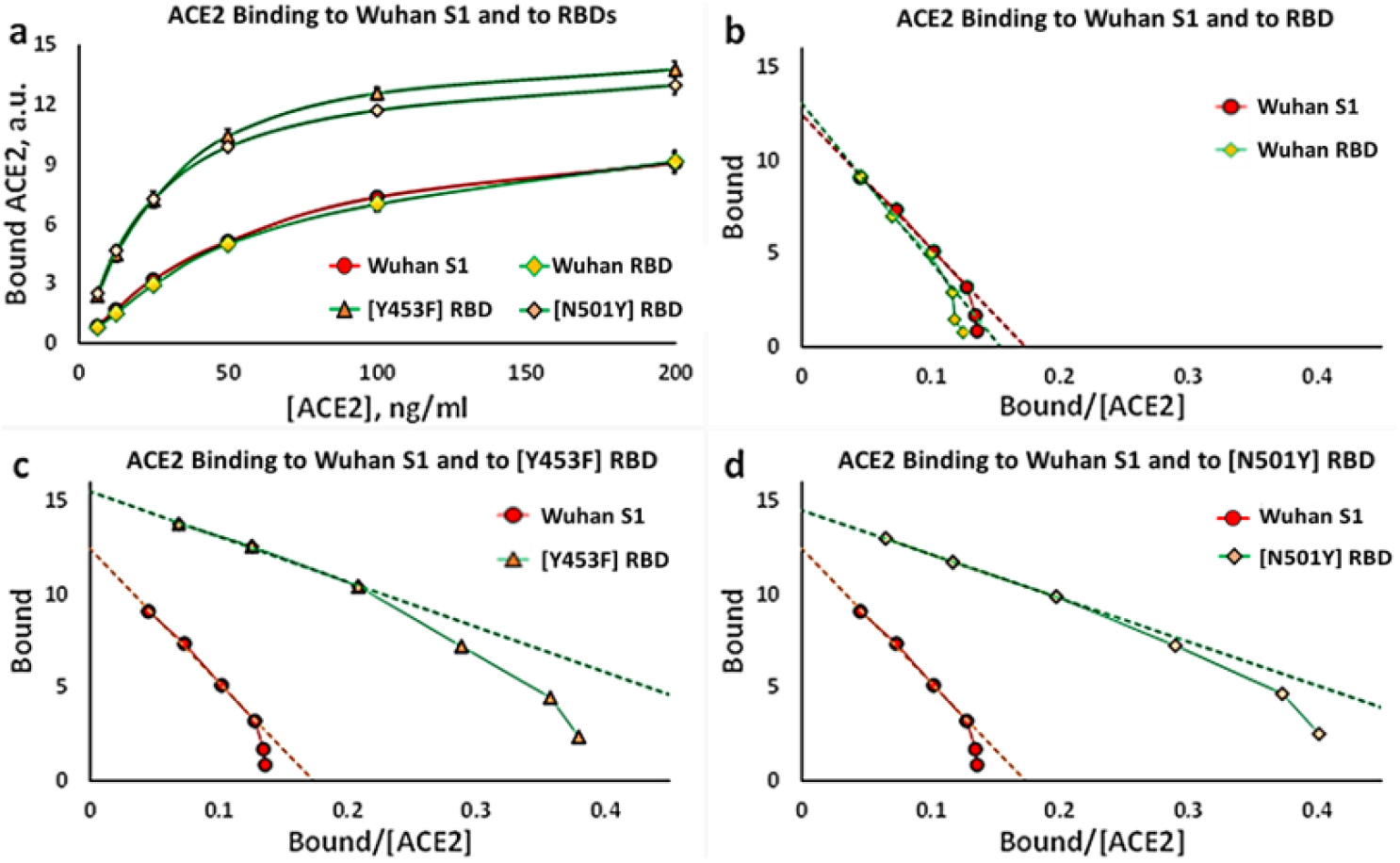
Binding kinetics for ACE2 to Wuhan and mutant spike variants. **(a)** Substrate-Binding plots for ACE2 binding to Wuhan RBD, Wuhan S1, and UK [N501Y] RBD proteins. (**b**) Eadie-Hoffstee plots for Wuhan RBD and Wuhan S1 proteins. (**c**) Eadie-Hoffstee plots for Wuhan S1 and Mink [Y453F] RBD proteins. (**d**) Eadie-Hoffstee plots for Wuhan S1 and UK [N501Y] RBD proteins. A *reduction* in slope corresponds to a *reduction* in K_D_, and thus an *increase* in affinity. Each point is the average ± SE for 5-6 independent experiments.

**Figure 3a** shows the substrate-binding plots for the recombinant S.Africa [E484K] RBD protein. Compared to the other RBD variants the S.Africa [E484K] RBD has a slightly more evident lag than the original Wuhan RBD. The ACE2 concentration region of 6.25-to-50 ng/ml is expanded in **Figure 3b** to emphasize this comparison with other variant RBD proteins. To further test for the ACE2 positive cooperativity mechanism we analyzed the S.Africa [E484K] RBD data using an Eadie-Hoffstee plot (**Figure 3c**). As for the other RBD variants the S.Africa [E484K] RBD protein has a profound “concave-down” structure for the lower values of ACE2 binding. The B_max_ is similar to that of the other variants, but the binding constant, K_D_, trends higher (**Table 1**). The Hill plot for the S.Africa [E484K] RBD protein is shown in **Figure 3d**. The slope for the S.Africa [E484K] variant, (n_H_), is 1.25 ± 0.05, and is significantly higher than for the other RBD proteins (**Supplementary Figure S5***). The elevated Hill constant is consistent with the relatively exaggerated concavity of the EH plot at low levels of ACE2 binding. Thus both the cooperativity and the affinity of ACE2 for the receptor binding domain can be modified by mutations not only in the RBD domain itself, but also by mutations elsewhere in the S1 region of the spike protein*.

**Figure 3.**
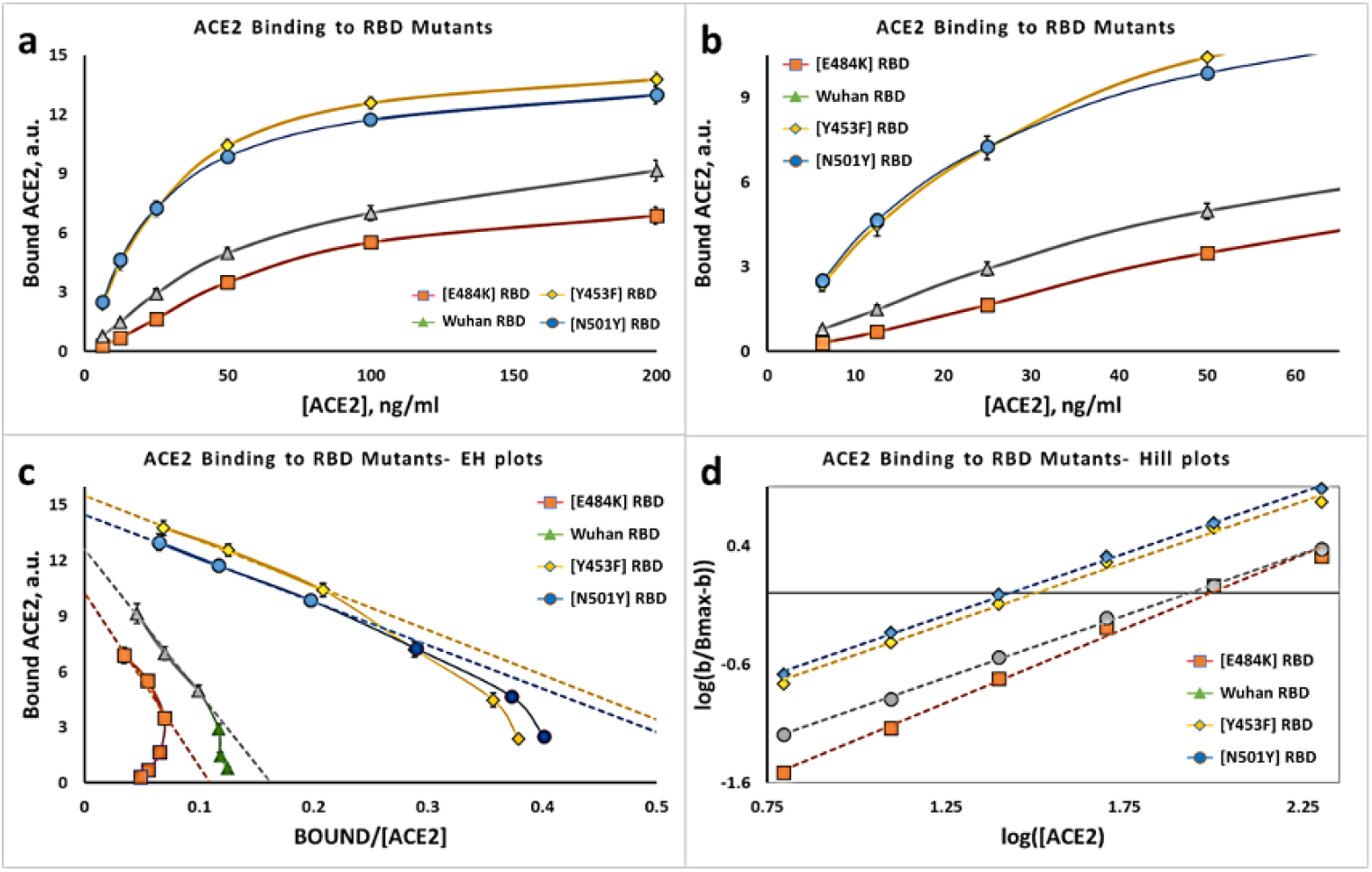
Binding kinetics for ACE2 to S.Africa [E484K] RBD, Mink [Y453F] RBD, UK [N501Y] RBD, and Wuhan RBD, **(a)** Substrate-Binding plots for ACE2 binding to RBD proteins. (**b**) Substrate - Binding plots for low ACE2 concentrations (**c**) Eadie-Hoffstee plots for mutant RBD proteins compard to Wuhan RBD. (**d**) Hill plots for mutant and Wuhan RBD proteins. Each point is the average ± SE for N= 5-6 independent experiments.

Sigmoid substrate-binding plots can also be due to autocatalysis, such as can occur during phase transitions. Therefore, to test for whether ACE2 binding to the Spike S1 derivatives might be due to autocatalysis as opposed to positive cooperativity, *per se*, we analyzed the ACE2 binding data using the recently reported Dhatt, Banerjee and Battacharrya (DBB) plot ^31^ (**Supplementary Figure S2).** In this plot pure Michaelis-Menten kinetics would be a straight horizontal line parallel to the X-axis. By contrast, autocatalysis would be a straight angular line, and positive cooperativity would be non-linear. The data show that the relationships are non-linear, thus corresponding to positive cooperativity. *Importantly, the data at any point define the apparent K*_*D*_, *and serve to show that the K*_*D*_ *values at higher concentrations of ACE2 are similar to values deduced from the linear portions of the Eadie-Hoffstee plots.*

### Cardiac glycoside drugs are competitive inhibitors of ACE binding to the RBD

To test for the ability of cardiac glycosides to inhibit ACE2 binding to the RBD protein, we measured ACE2 binding to the RBD protein in the presence of digitoxin, digoxin and ouabain (**Figure 4a**). Reduced ACE2 binding in the presence of these drugs is evident. The substrate-binding isotherms also indicate that the sigmoid appearance is sustained. Thus the positively cooperative binding mechanism is still present. *The DBB plots also provide evidence for a sustained positive cooperativity mechanism for ACE2:RBD binding in the presence of the cardiac glycoside drugs* (see **Supplemental Figure S3).**

**Figure 4.**
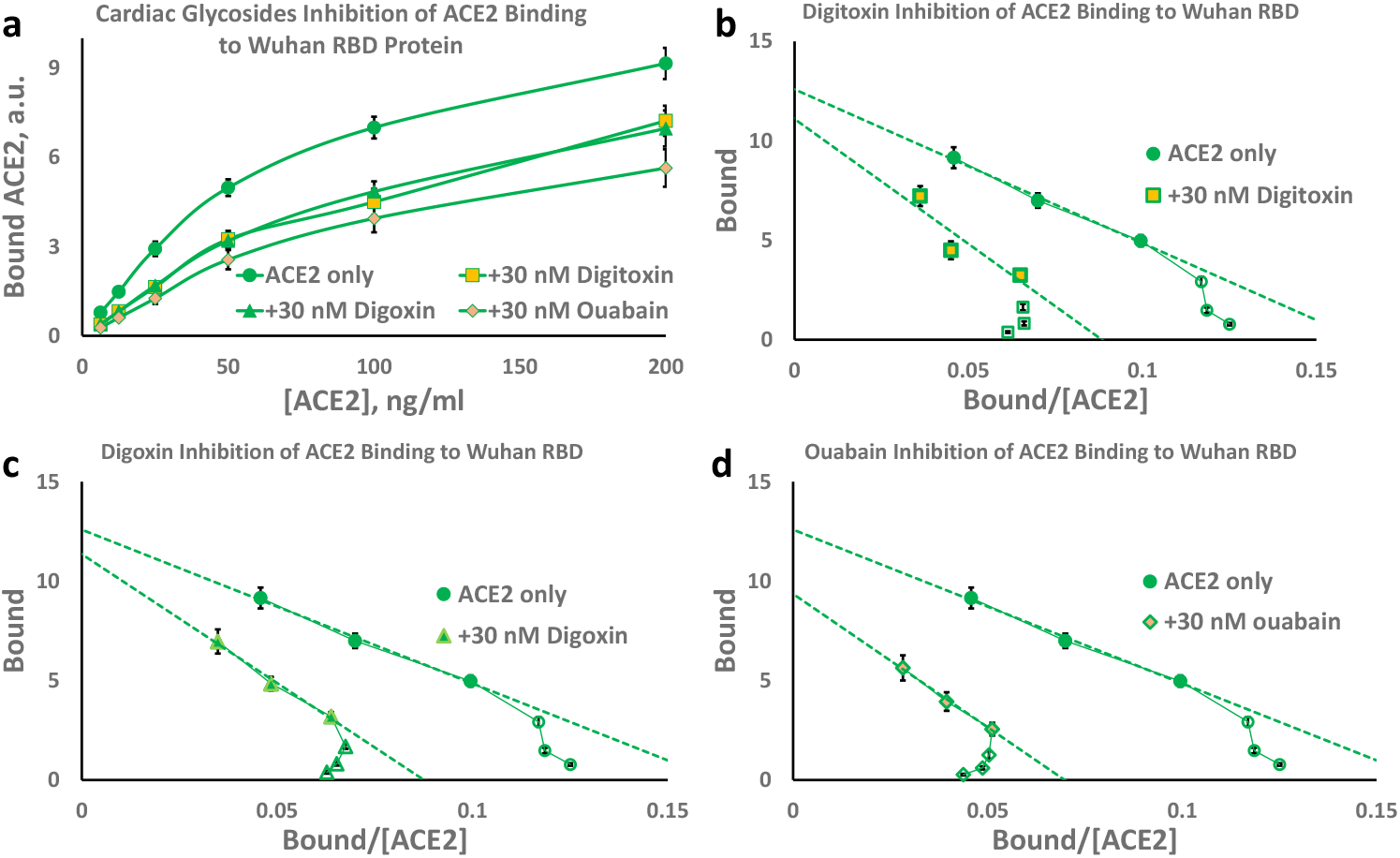
Inhibition of ACE binding to the Wuhan RBD by cardiac glycoside drugs. **(a)** Substrate-Binding plots for inhibition by digitoxin, digoxin and ouabain (30nM). (**b**) Eadie-Hoffstee (EH) plots for digitoxin inhibition. (**c**) EH plots for digoxin inhibition. (**d**) EH plots for ouabain inhibition. Each point is the average ± SE for 5-6 independent experiments.

In addition, the Eadie-Hoffstee plots for all three drugs show that the cardiac glycosides differentially increase the K_D_ values while keeping the B_max_ values approximately constant (see **Figure 4b, c** & **d**). Similar data were obtained for the Wuhan [D614] S1 protein (**Figure 5**) and for the European mutant [D614G] S1 protein (**Figure 6**). These data are consistent with a competitive inhibition mechanism. The individual EH plots show that these drugs all appear to lower the affinity for ACE2 by 50-100% while keeping the B_max_ approximately constant. *These kinetic data are summarized in **Table 2**, and indicate that of the three drugs tested ouabain is the most potent inhibitor, followed by digitoxin and digoxin.*

**Figure 5.**
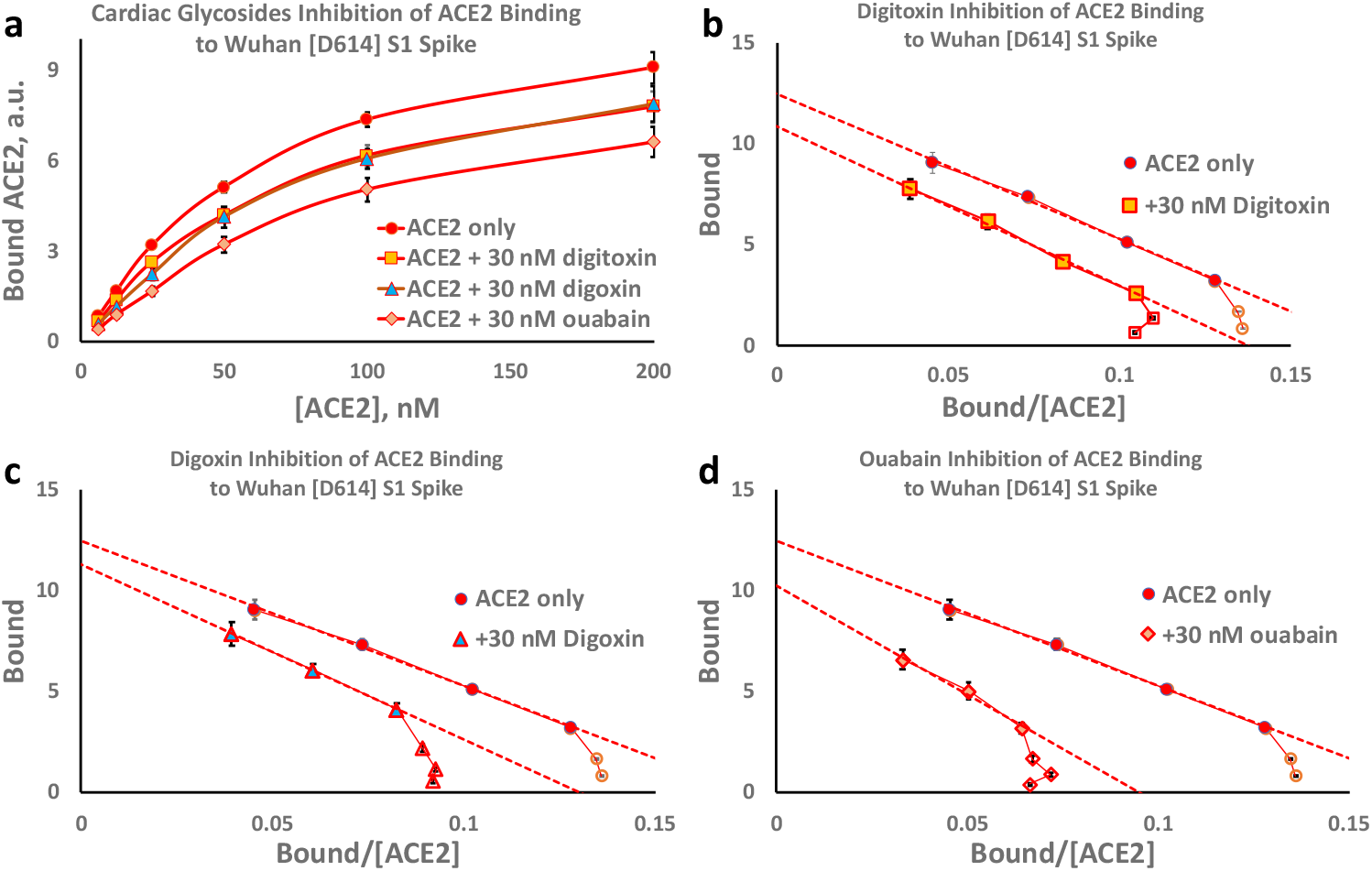
Inhibition of ACE binding to the Wuhan [D614] spike S1 by cardiac glycoside drugs. **(a)** Substrate-Binding plots for inhibition by digitoxin, digoxin and ouabain (30nM). (**b**) Eadie-Hoffstee (EH) plots for digitoxin inhibition. (**c**) EH plots for digoxin inhibition. (**d**) EH plots for ouabain inhibition. Each point is the average ± SE for 5-6 independent experiments.

**Figure 6.**
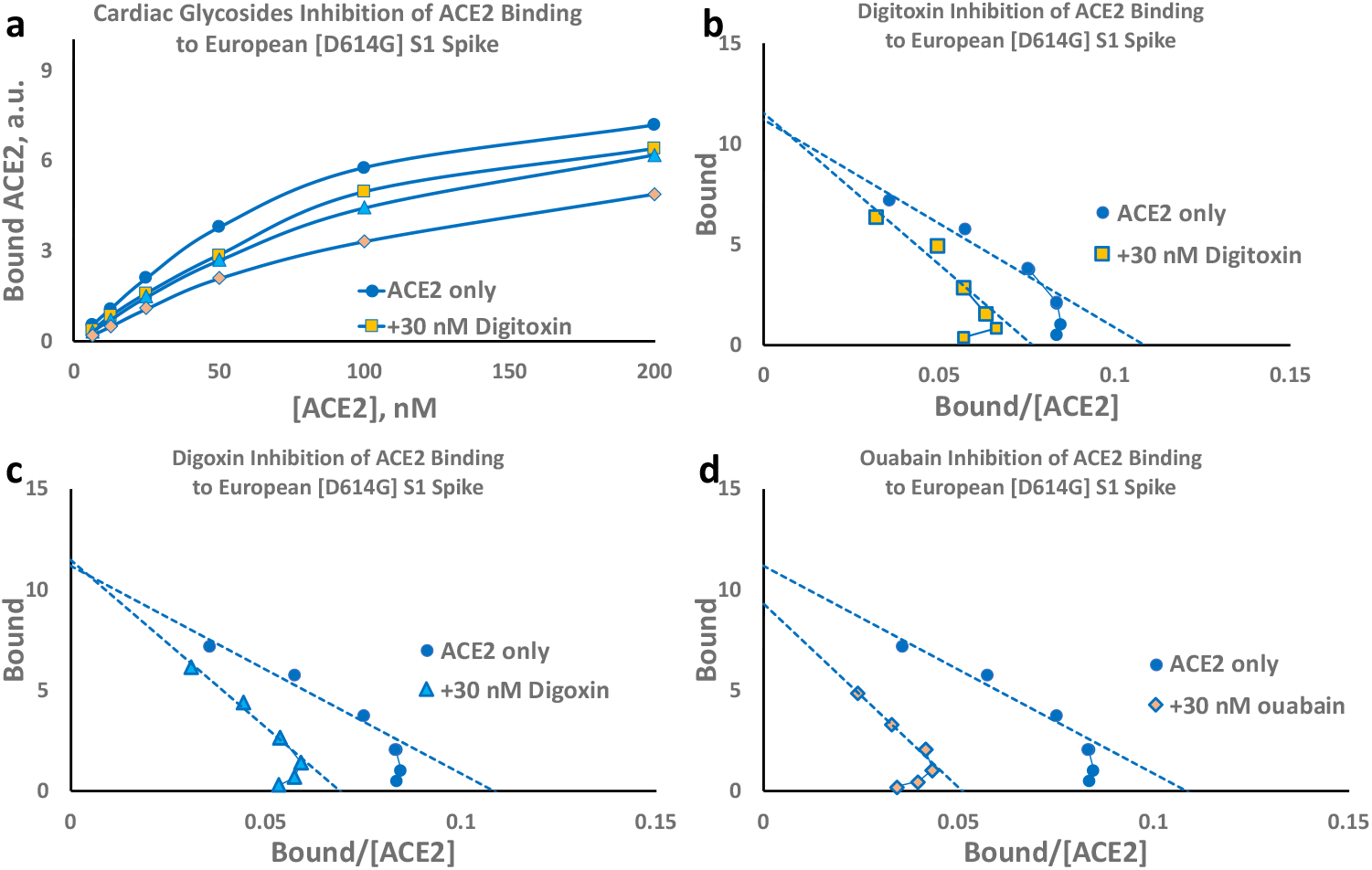
Inhibition of ACE2 binding to the European [D614G] spike S1 by cardiac glycoside drugs. **(a)** Substrate-Binding plots for inhibition by digitoxin, digoxin and ouabain (30nM). (**b**) Eadie-Hoffstee (EH) plots for digitoxin inhibition. (**c**) EH plots for digoxin inhibition. (**d**) EH plots for ouabain inhibition. Each point is the average ± SE for N = 5-6 independent experiments.

**Table 2.** Inhibition constants Ki for Cardiac Glycoside Inhibition of ACE2 Binding to SARS-CoV-2 Spike Variants. Values are based on calculating Ki values for all concentrations of ACE2 and averaging all values for the individual recombinant proteins. Each point is the average ± SE for 5-6 independent experiments.

| | Inhibitor $K_i$, nM ( ± SE ) | | | | |
| --- | --- | --- | --- | --- | --- |
| Spike Variant | Digitoxin | Digitoxigenin | Digoxin | Digoxigenin | Ouabain |
| Wuhan [D614] S1 | 69 ( ± 8 ) | 63 ( ± 6 ) | 53 ( ± 6 ) | 35 ( ± 3 ) | 24 ( ± 2 ) |
| European [D614G] S1 | 71 ( ± 13 ) | 54 ( ± 5 ) | 48 ( ± 5 ) | 34 ( ± 4 ) | 21 ( ± 2 ) |
| Wuhan RBD | 28 ( ± 3 ) | 27 ( ± 2 ) | 29 ( ± 2 ) | 21 ( ± 2 ) | 16 ( ± 1 ) |
| S.Africa [E484K] RBD | 43 ( ± 13 ) | 60 ( ± 33 ) | 62 ( ± 23 ) | 30 ( ± 4 ) | 21 ( ± 4 ) |
| Mink [Y453F] RBD | 71 ( ± 12 ) | 41 ( ± 4 ) | 67 ( ± 12 ) | 26 ( ± 2 ) | 45 ( ± 5 ) |
| UK [N501Y] RBD | 62 ( ± 11 ) | 39 ( ± 5 ) | 43 ( ± 5 ) | 23 ( ± 2 ) | 37 ( ± 4 ) |

\*: Each value is calculated from N = 5-6 independent experiments, ± SE

Digitoxin and digoxin have three sugars on their 3’-OH groups, while the more potent inhibitor ouabain has only one sugar (see **Supplemental Figure S1b**). We therefore tested whether the sugar moieties on digitoxin or digoxin contributed to inhibitory activity. Digitoxingenin and digoxigenin are the sugar-free analogues of digitoxin and digoxin, respectively. We found that for ACE2 binding to the Wuhan S1 spike protein, 30 nM digitoxigenin was approximately as inhibitory as digitoxin **(Supplemental Figure S4a).** A similar result was noted for digoxin and digoxigenin (**Supplemental Figure S4b**). We also found similar results for inhibition of ACE2 binding to the European mutant [D614G]S1 (**Supplemental Figures S4c and S4d**), and to the Wuhan RBD protein (**Supplemental Figures S4e** and **S4f**). The DBB plots in **Supplemental Figure S3** also show that the sugar-free forms of digitoxin and digoxin are very similar inhibitors to their respective parental drugs, and that the positive cooperative behavior of the ACE2 binding isotherm is preserved. Additionally, **Supplementary Figure S4** indicates that the blocking mechanism for both sugar-free drugs is by competitive inhibition. The K_i_ values for digitoxigenin and digoxigenin are included in **Table 2**. *The data therefore suggest that the pharmacophore for digitoxin and digoxin inhibition of ACE2 binding to the RBD may reside in their respective steroid nuclei.*

Finally, we tested the ability of cardiac glycoside drugs to inhibit ACE2 binding to the S.Africa [E484K] RBD protein, the UK [N501Y] protein and the Mink [Y453F] RBD protein. **Figure 7a** shows the Substrate-Binding plots for ACE2 binding to the S.Africa [E484K] RBD protein. The inhibitory effects are clearly apparent for the cardiac glycoside drugs ouabain, digitoxin and digoxin, and the two sugar-free derivatives, digitoxigenin and digoxigenin. The sigmoid character of the ACE2 binding isotherm is also evident, and is more clearly shown in **Figure 7b** for the low ACE2 concentrations. **Figure 7c** shows the Eadie-Hoffstee plots for both drugs and derivatives. All are concave-down at low concentrations of ACE2. Thus the apparent positive cooperativity for ACE2 binding is preserved in the presence of all three cardiac glycosides and the two sugar-free derivatives. Furthermore, all of these drugs and their sugar-free derivatives have inhibitory activity, although the most active drug is ouabain. Finally, the B_max_ values for these drugs are similar while the slopes of the linear portion of the EH plots at high ACE2 concentration are systematically higher than the control. Thus the inhibitory mechanisms appear classically competitive. **Figure 7d** shows the Hill plots for all five cardiac glycosides and sugar-free derivatives. Equivalent results for the Mink [Y453F] RBD protein and the UK [N501Y] RBD protein are shown in **Supplementary Figures S6** and **S7** respectively. The calculated Ki values for these drugs and their sugar-free derivatives are shown in **Table 2**. Ouabain has the lowest Ki value of the three cardiac glycoside drugs. Digoxigenin has a slightly lower Ki value for Mink [Y453F] and UK [N501Y]. *Thus relative cardiac glycoside potency for inhibition of ACE2:RBD binding appears to depend both on the drug structure, and on the mutation.*

**Figure 7.**
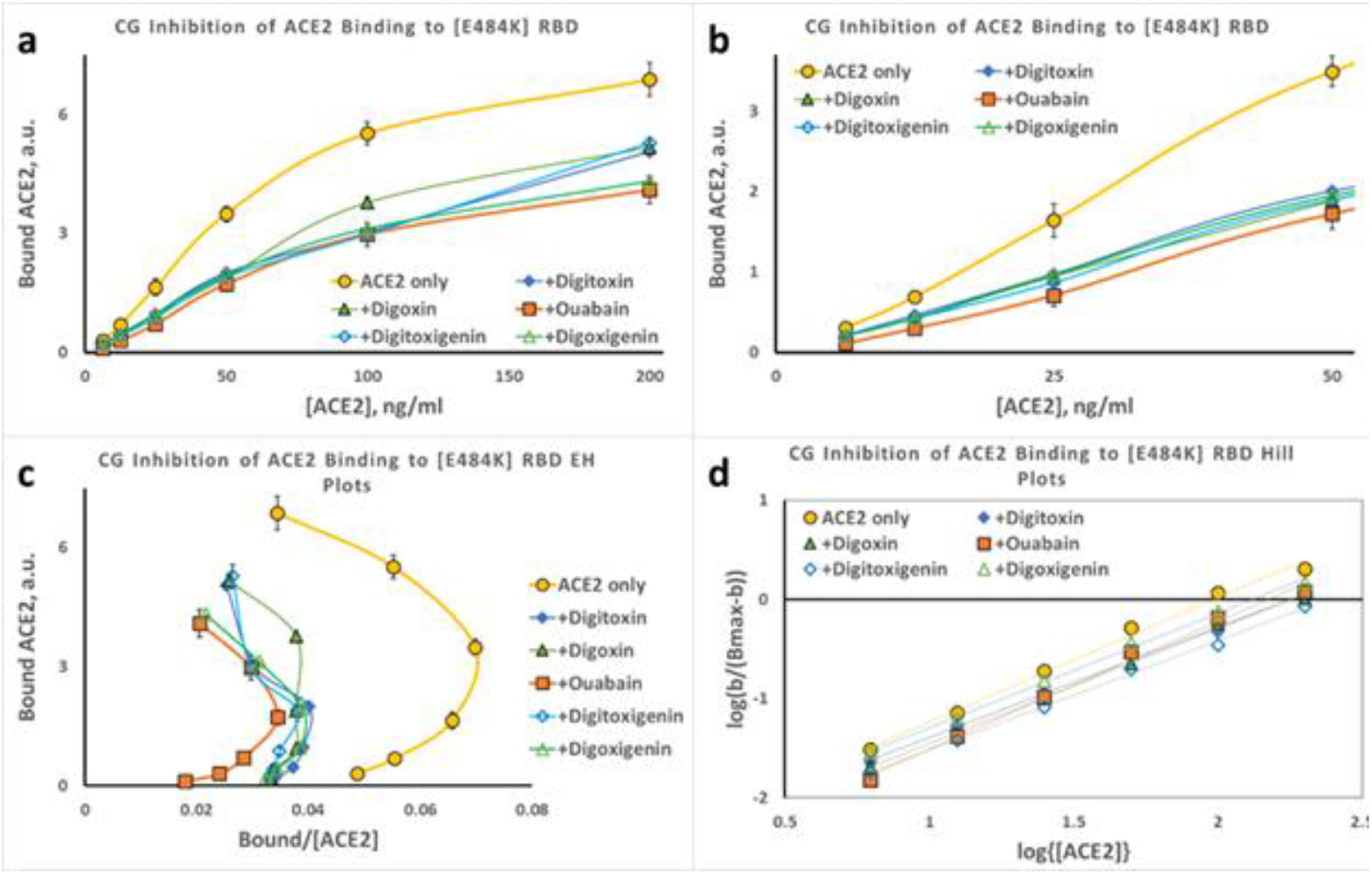
Inhibition of ACE2 binding to the S.Africa [E484K] RBD by cardiac glycoside drugs. **(a)** Substrate-Binding plots for inhibition of ACE2 binding to S.Africa [E484K] RBD protein by digitoxin, digoxin, ouabain, digitoxigenin and digoxigenin (30nM). (**b**) Data from (**a**) at low concentrations of ACE2. (**c**) Eadie-Hoffstee (EH) plots of data in part (**a**). Hill plots for each titration from part (**a**). Magnitude and error for Hill coefficient (n_H_) given in **Supplemental Figure 5**. Each point is the average ± SE for N = 5-6 independent experiments.

The Hill plot slopes from **Figure 7d** and **Supplemental Figures S6d** and **S7d** vary between ~1.1 and ~1.3 for all tested variant Spike proteins, and are summarized in **Supplementary Figure S5a**. The highest values are for S.Africa [E484K], regardless of which drug or drug-derivative is present in the analysis. To determine the significance of these observed differences we calculated p-values for the difference between the observed n_H_ value for each condition and n_H_ =1.0, where cooperativity is zero (**Supplemental Figure S5b**). The mutant with the highest significance, independent of the treatment, is the S.Africa [E484K]. The drug that most significantly affects the most mutants is ouabain. *Operationally, a high value of n_H_ is associated with a more lengthy lag in ACE2 binding at low concentrations of ACE2*.

### Cardiac glycosides block penetration of Spike-pseudotyped VSV virus into lung cells

To test whether cardiac glycosides blocked the viral penetration process we tested drug effects on cell entry by luciferase (*luc*)-loaded Spike-pseudotyped VSV(S) viral particles. To further focus the experiment on the earliest entry steps we also used a triple tandem time-of-addition method. Briefly, VSV(S) particles were *first* preincubated for one hour with cardiac glycoside drugs in DMEM at 37°C. *Secondly*, the particles and drugs were then preincubated with the A549 cells for two hours at 37°C. *Thirdly*, the viral particles and drugs were washed away and replaced by complete DMEM medium. The A549 cells were then incubated for 24 hours at 37°C before assay for luciferase. **Figure 8a** shows the results of luminescence background-corrected activity of *luc*-loaded pseudotyped VSV(S) over the active ranges for digitoxin, digoxin and ouabain. Ouabain was found to potently suppress viral entry. Digitoxin was less potent and digoxin was the least potent. Based on these data the Hill plot-based EC-50 value for ouabain was calculated to be 16 ± 1 nM (N=3, ± SE). The positive and negative control values are given in **Supplemental Table S1**. The inhibition data are similar when measured either by ratio to the luminescence control (**Figure 8b**), or by % of positive control (**Figure 8c**). As a negative control, and to test for whether human ACE2 receptor was required for cell penetration, mouse 3T3 cells were substituted for human A549 cells. Mouse 3T3 cells were found to be inactive (**Figure 8d**). Finally, as a positive human control, post convalescent serum from an unidentified, recovered COVID-19 patient was found to inhibit penetration of the SARS-CoV-2 spike pseudotyped VSV(S) in a dose-dependent manner (**Supplemental Figure S8).** *Since the SARS-CoV-2 spike pseudotyped VSV(S) only reports on viral penetration, and is not infectious, the data indicate that ouabain most potently blocks viral penetration into human lung cells, likely at a very early stage of the entry process.*

**Figure 8.**
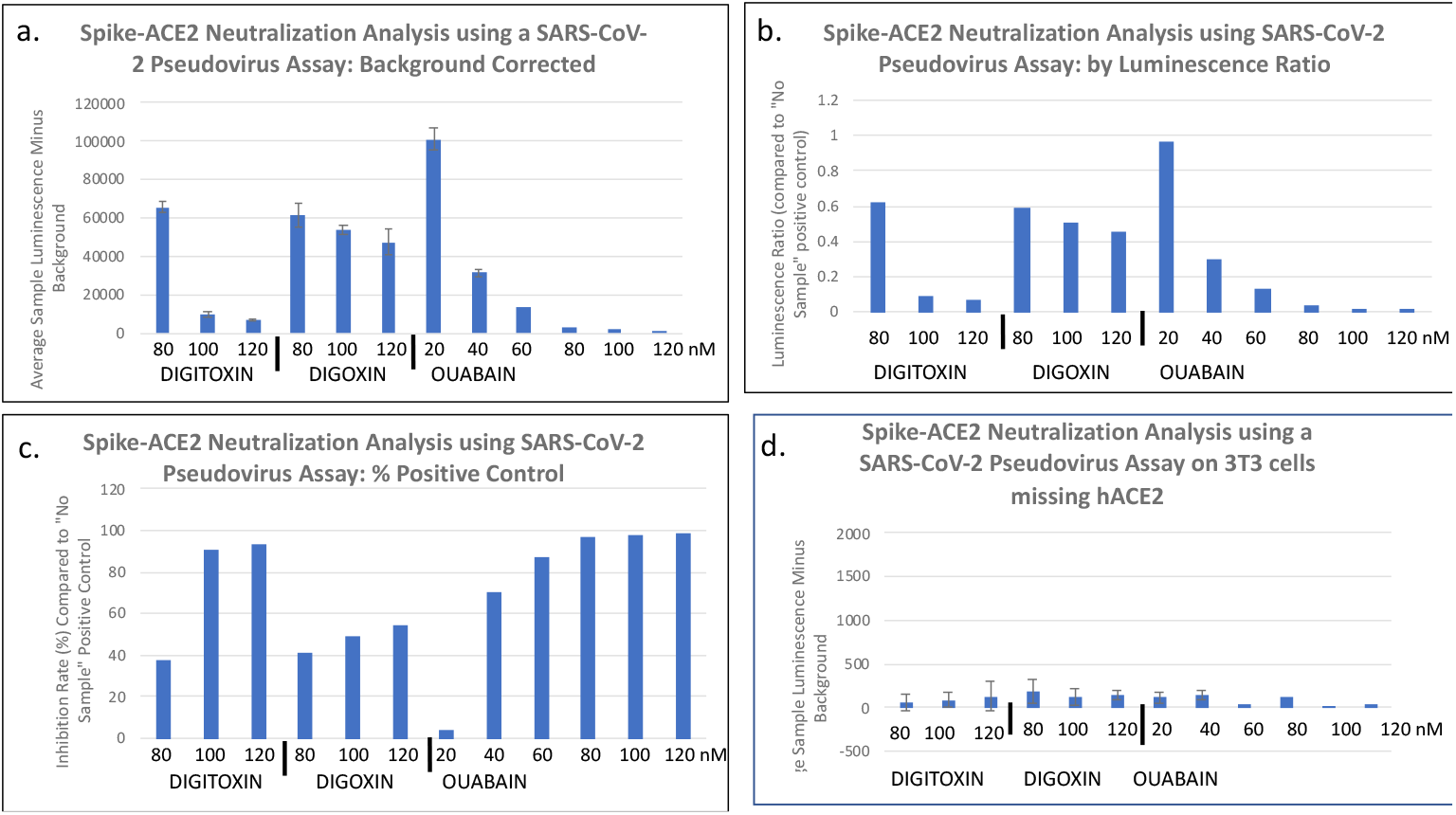
Inhibition of pseudotyped VSV(S) penetration into A549 lung cell by cardiac glycosides. **(a)**Measurement of inhibition by luminescence background subtract. VSV(S) is pre-treated with drugs for 1 hr, and then exposed to cells in DMEM for 2 hours. VSV(S) mixture is then replaced by whole medium and incubated for 24 hours. Data are the averages of 3 independent experiments. Error bars are ± SE. Significance based on p <.05. (**b**) Measurement by luminescence ratio. (**c**) Measurement by % of positive control. (**d**) Negative control with mouse 3T3 cells missing hACE2, measured by luminescence background subtract method.

## Discussion

For the SARS-CoV-2 virus to infect a target cell, the Receptor Binding Domain on the viral Spike protein must bind to the receptor protein ACE2 on the cell surface. However, fully understanding the factors controlling this binding process is only at an early stage. Here we have tested the hypothesis that cardiac glycoside drugs such as ouabain, digitoxin, and digoxin might block the binding reaction between ACE2 and the Spike (S) protein, and thus block viral penetration into human lung cells. In this paper both parts of this hypothesis have been experimentally validated in three overlapping tests. Firstly, we found that ACE2 binding to various spike variants occurred by a positively cooperative mechanism. Not unexpectedly, we also found that mutations in the RBD, specifically Mink [Y453F] and UK [N501Y], significantly *increased* ACE2 binding affinity, while the S.Africa [E484K] mutation trended towards *reduction* in ACE2 binding affinity. We also validated a previous finding that the [D614G] mutation, on a part of the spike S1 protein *outside* of the RBD, also *reduced* ACE2 binding affinity to the RBD. *Thus the effect of a mutation that enhances infectivity need not necessarily increase the affinity of ACE2 for the RBD*. In addition, as hypothesized we found that ouabain, digitoxin and digoxin were also high-affinity competitive inhibitors of ACE2:Spike binding. In addition, we found that the *relative inhibitory potencies of these drugs, as defined by the K*_*i*_, *were mutation-and drug-dependent.* Lastly, we found that ouabain and digitoxin, but not digoxin, potently blocked SARS-CoV-2 Spike pseudotyped virus penetration into human lung cells. Inasmuch these drugs are widely available, and are clinically safe for those with normal hearts ^29^, it is quite possible that these common cardiac medications could be repurposed for anti-COVID-19 therapy.

In retrospect the positively cooperative mechanism by which ACE2 binds to the spike might have been anticipated. For example it has been recently reported that when either the RBD protein or the intact spike protein binds to ACE2 there is a 3-10-fold increase in the intrinsic ACE2 carboxypeptidase activity ^32^. Thus functional changes in ACE2 can be induced by interaction with spike proteins. Furthermore, structural analysis by Cryo-EM shows that the binding of RBD to ACE2 induces a ~12° movement of the ACE2 C-terminal domain toward the N-terminal domain, and also induces an opening of the ACE2 substrate binding pocket ^33^. Such changes in conformation are intrinsic to the two major models for positive cooperativity, the “induced fit” model ^30,34,35^ and the earlier “sequential” or “symmetry” model ^30,36^. Furthermore, the modest Hill coefficients shown here for ACE2:Spike interactions are also consistent with enzymatic and biophysical data previously reported for positively cooperative monomeric systems such as glucokinase ^37^ and trypsin-like proteases ^38^. These data therefore might be consistent with an “induced fit” hypothesis, in which ACE2 could initially bind to the RBD with low affinity, and then undergo further conformational changes to accomplish a higher affinity interaction. On the other hand, dimers of native ACE2 have been shown to be preferred as a binding partner for the RBD for a pseudotyped SARS-CoV-2 spike ^33^. Thus monomers of ACE2 bind poorly to intact viral Spikes, compared to the binding by ACE2 dimers. In addition, it has been reported that more than one RBD in a triple protomer spike can elevate into the open state and bind to multiple host ACE2 extracellular domains ^39,40^. Finally, recent *in silico* modeling has suggested that higher concentrations of recombinant extracellular ACE2 might trend towards a dimeric polymerization state ^41^. *We conclude that further analysis will be necessary to distinguish whether one or both models might best describe the experimental data.*

Based on data shown here, we have also shown that other regions in the Spike complex might have conformational effects on the RBD and thus its affinity for ACE2. The [D614G] mutation in the S1 protein may be an example of how changes at some distance from the RBD sequence can raise the K_D_ for ACE2 binding to the RBD. How this happens, and why there are possible advantages to the virus for infectivity, are insights yet to be discovered. However, it has been reported that the [D614G] mutation increases spike flexibility, and spike density by 4-5 fold ^14,15^. Thus in exchange for a reduction in the affinity for ACE2, other properties that enhance infectivity may have been acquired*. Nonetheless, the inhibition constants for the cardiac glycosides were not significantly affected by the [D614G] mutation.*

The anti-viral properties of cardiac glycosides have been well documented, not only for some RNA viruses such as coronaviruses but for some DNA viruses as well ^42^. In the case of RNA viruses, ouabain and digoxin have been reported to block native MERS-CoV entry into green monkey kidney Vero cells ^23^. Digitoxin has also been reported to have the same effect on MERS-CoV entry into Vero cells ^24^. Recent time-of-addition studies on native SARS-CoV-2 with Vero cells have also been interpreted to indicate that ouabain may block viral entry, with an EC_50_ of 24 nM 22. However, digoxin, with an EC_50_ value of 43 nM, did not appear to block viral entry, and has been hypothesized to interfere with infection at some intracellular site ^22^. Consistently, the analysis reported here directly shows that ouabain and digitoxin, but less so digoxin, act as blockers of viral entry into human lung cells. In sharp contrast, the effects of ouabain and other cardiac glycosides on other RNA and DNA viruses have *not* been associated with viral entry. Rather the effects have been generally on downstream activities associated with virus propagation and infection ^42^. Thus the specific anti-viral activity of blocking viral entry into cells by low concentrations of ouabain, and to a lesser extent by digitoxin, appears to be limited to the coronaviruses, and here specifically to SARS-CoV-2. *Therefore we suggest that the high affinity and mutation-dependent specificity of the ACE2:RBD binding reaction may point to a biologically important process that might be used to therapeutic advantage for COVID-19.*

Cardiac glycosides such as digitoxin have a recent history as potent inhibitors of the TNFα-dependent proinflammatory *host response* to influenza virus infection, also known as cytokine storm ^43^. Consistently, digitoxin has also been reported to be a potent and efficacious suppressor of TNFα-dependent proinflammatory cytokine expression in cystic fibrosis cells and patients as well ^28,44^. Digitoxin also inhibits tumor growth and NFκB signaling in a pre-clinical rat model of castration resistant prostate cancer ^45^. Recently, digitoxin was reported to be among the top ten inhibitors of TNFα-activated NFκB signaling in a screen of 2,800 FDA approved drugs ^46^. The mechanism for digitoxin inhibition of TNFα signaling to NFκB is due to blocking the interaction between the proinflammatory TNFα/TNFR1 complex and the TNFα Associated Death Domain (TRADD) protein ^47^. TRADD is the first intracellular adaptor for the activated TNFα/TNFR1 complex. Mutant CFTR also fails to suppress TRADD expression, thus explaining why cystic fibrosis patients present with chronic TNFα-dependent proinflammatory disease in lung and other organs ^48^. By contrast, digitoxigenin and digoxigenin, the sugar-free analogues of digitoxin and digoxin, respectively, are relatively inactive as blockers of the TNFα-dependent host response ^49^. Consequently, the order of suppression of host-specific proinflammatory suppression potency is [digitoxin > ouabain >> digoxin >>> digitoxigenin, digoxigenin]. This is a different potency order for inhibition of ACE2:Spike binding. *We conclude that different mechanisms must mediate cardiac glycoside-dependent inhibition of TNFα-dependent host defense and inhibition of SARS-COV-2 entry into target cells*.

Finally, safety is an issue to be considered, even when treating a lethal disease like COVID-19. As a class, the cardiac glycosides such as digitoxin, digoxin and ouabain are said to have a narrow therapeutic index when treating a patient with heart failure ^29^. A narrow therapeutic index means that medicinal dose and the toxic dose are very close, and care must be taken to avoid toxicity. However, according to Goodman and Gilman, the authoritative pharmacology text, “…If subjects with normal hearts ingest large but not lethal quantities of digitalis, either in an attempt at suicide or by accident, premature impulses and rapid arrhythmias are infrequent.” ^29^. Digitalis is an extract of *Digitalis* plant species containing a mixture of digitoxin and digoxin. In support of this recommendation, *normal* subjects have been studied quantitatively after treatment with purified ouabain, digitoxin and digoxin. No toxicity or adverse events have been reported. In specific examples with ouabain Mason and Braunwald ^50^ injected 0.50-0.60 mg ouabain into 12 normal subjects, aged 18-49, and measured forearm vascular resistance and venous tone. No adverse effects were reported. Selden and Smith ^27^ administered 0.25 mg ouabain to normal human volunteers, in order to determine ouabain pharmacokinetics. The peak plasma concentration was found to be 12 ng/ml, or 20 nM, with a half-life of 19-24 hours. This peak value was slightly more than the EC_50_ of 16 nM measured here for inhibition of virus penetration into human lung cells. These volunteers were also treated daily for 9 days with 0.25 mg ouabain, reaching a plateau in 4-5 days. No adverse events were reported. More recently, *digitoxin* has been administered by mouth to 16 patients with cystic fibrosis (CF), a proinflammatory genetic lung disease ^28,44^. Subjects were administered 0.05 or 0.1 mg/day for 28 days in a double blind, placebo controlled study. Proinflammatory signaling in airway epithelial cells was significantly suppressed at the higher dose, and again no adverse events were associated with the study drug. The peak serum digitoxin concentration in those CF patients who were treated with 0.1 mg/day reached 10 ng/ml, or 13 nM, by day 28. In addition, over the years, multiple studies on digoxin in cardiovascularly normal subjects have been reported ^51–54^. In one instance 0.25 mg *digoxin* was administered to cystic fibrosis patients, daily for one week, with no adverse events associated with the study drug ^51^. Finally, in a study of 47,884 men, those taking digoxin for heart failure for >10 years had 25% lower prostate cancer risk (p < 0.001) ^55^ Thus toxicity does not necessarily accompany prolonged treatment with these drugs, even for those with cardiac disease. Lastly, and just in case toxicity does occur in cardiac patients or others, Digibind^R^ is now routinely available in poison control centers. Digibind ^R^ is an antibody preparation against the sugar-free steroid nucleus common to cardiac glycosides.

It is a limitation of our study that we have not tested whether changes in spike conformation in response to ACE2 binding might also be occurring. Certainly infectivity enhancing mutations such as Mink [Y453F], UK [N501Y] and S.Africa [E484K] profoundly affect ACE2 binding kinetics. But whether RBD conformation intrinsically changes in response to ACE2 binding is a study for the future. An additional limitation of our study is that we have utilized a *luc*-loaded, SARS-CoV-2 pseudotyped virus as a test for biological activity rather than a native SARS-CoV-2 virus. However, it was only with a viral particle model that we could unambiguously demonstrate that cardiac glycosides inhibited viral entry. As mentioned above, ouabain itself has also been previously demonstrated to block infectivity by native SARS-CoV-2 virus, albeit in a green monkey Vero cell ^22^.

### Conclusion

Ouabain and digitoxin competitively inhibit ACE2 binding to the SARS-CoV-2 Receptor Binding Domain (RBD), and also block virus penetration into human lung cells. Clinical concentrations of cardiac glycosides are relatively safe for subjects with normal hearts, and it is therefore possible that these drugs could be repurposed for COVID-19 therapy.

## Materials and Methods

### Chemicals and biologics

Digitoxin, digoxin, ouabain, digitoxigenin and digoxigenin were purchased from Sigma-Aldrich. Drugs were solubilized as 4 mM stock solutions in either 95% ethanol or 100% DMSO, and aliquots were serially diluted in the same solvent, and then into assay medium. In assays containing ethanol or DMSO, the final solvent concentrations were 0.01% or less. Recombinant proteins produced in human T293 cells were obtained as follows: Recombinant Human Angiotensin Converting Enzyme 2 (ACE2) (Cat # 230-30165; lot 04U06020TW) was purchased from RayBiotech (Peachtree Corners, GA, 30092). Human ACE2 Biotinylated Antibody (Cat # BAF933) and Recombinant Spike S1 RBD protein (Cat # NBP2-90982) were obtained from R&D Systems (Minneapolis, MN, 55413) and NOVUS Biologicals (Centennial, CO, 90982), respectively. Recombinant SARS-CoV-2 Wuhan RBD protein (Cat # 40592-VO8H), Spike [D614] S1 protein (Cat # 40591-V02H), [D614G] S1 protein (Cat # 40591-V08H3), Mink [Y453F] RBD protein (Cat # 40592-V08H80), UK [N501Y] RBD (Cat #40592-V08H82) and S. Africa (E484K) RBD proteins (Cat # 40592-Vo8H84) were purchased from SinoBiologicals U.S. Inc. (Chesterbrook, PA 19087).

### Enzyme Linked Immunosorbant Assay (ELISA) for interaction between Spike proteins and ACE2

Purified recombinant SARS-CoV-2 Spike proteins were individually dissolved in coating buffer (16 mM Na_2_CO_3_, 34 mM NaHCO_3_, pH 9.6) at a concentration of 2 μg/ml Spike protein. This solution, in 100 μL aliquots, was then added to wells of Costar 96 well plates (Corning, Corning, NY; Catalog # 2592) and incubated overnight at 4°C. The next day, wells were washed 3 times in Phosphate Buffered Saline with 0.05% Tween 20 (PBST) at room temperature. Wells were then blocked with 300 μL Blocking Buffer (0.5% Bovine Serum Albumin (BSA), dissolved in PBST) for two hours at 37°C. Wells were then washed 3 times in PBST at room temperature (68°F). When cardiac glycosides were to be added, they were dissolved in Reagent Diluent (0.5% BSA in PBST), added in 100 μL volumes to each well, and incubated at 37°C overnight. The next day, plates were inverted gently on paper towels for 5 minutes. A volume of 100μL dilutions of ACE2 were added to each well, supplemented with cardiac glycosides as appropriate, and incubated at 37°C for two hours. Wells were then washed 3 times with PBST at room temperature. A working concentration of 1 μg/ml human ACE2 biotinylated antibody (R&D Systems, Minneapolis, MN) was prepared, and 100 μL of this solution was added to each well. The plates were protected from light, and incubated at 37°C for one hour. Wells were then washed three times in PBST. HRP-Strepavidin (R&D Systems, Minneapolis, MN) was diluted 1:200, and 100 μL of the solution was added. Plates were then incubated at 37°C for one hour, and were washed three times in PBST. A substrate solution was prepared from R & D Systems kit components Solution A and Solution B, mixed 1:1, and 100 μL added to each well. Plates were incubated in total darkness for 15 minutes. A volume of 50 μL Stop Solution (2N H_2_SO_4_) was then added to each well. The wells were then read at 450 nm and 570 nm on a FLUOstar -Optima scanner (BMG Labtech, GmbH, Ortenberg, Germany). Individual experiments consisted of 3 or 4 technical replicates for each condition. A non-specific control without spike protein was always included in every assay. Except as noted, each experiment was independently repeated 5-7 times. The final results are given as averages ± Standard Errors of all independent experiments.

### Cells and pseudotyped virus

SARS-CoV-2 Spike pseudotyped Vesicular Stomatitis Virus (VSV(S)) particles were generated and tested using pseudovirus rVSV-ΔG-pCAGGS/Spike-luciferase) ^56–59^ in RayBiotech laboratories (RayBiotech, Peachtree Corners, GA, 30092). Briefly, VSV(S) particles were preincubated in different concentrations of cardiac glycosides in DMEM for 1 hour at 37°C. Thereafter, 100 μL of the mixture was transferred to a semi-confluent culture of A549 human alveolar basal epithelial cells growing at a density of 20,000 cells/well in a 96-well white cell culture plate with a flat clear bottom. Two hours later, VSV(S) particles were removed and the cells were washed 3 times in DMEM. The medium was then replaced with DMEM with complete growth medium containing 10% FBS and 2% penicillin-streptomycin, and the cells were incubated for 24 hours at 37oC/ 5% CO_2_. To measure luciferase, 50 μL of luciferase substrate replaced the medium in each well. The plate was shaken for 3 minutes at 300 rpm to lyse cells and equilibrate samples, and luminescence was then measured.

### Statistics

To determine the significance of changes in kinetic parameters of ACE2 binding to SARS-CoV-2 Spike or RBD mutants we have applied least-squares regression to the linearized Eadie-Hoffstee plot of the binding data, and determined the statistical significance of the difference between the slopes (*i.e*., K_D_’s) and between the intercepts (*i.e*., B_max_) using R or Stata statistical packages. K_i_ values were calculated for each data point in the linear range depending on the inhibition mechanism. For pseudovirus experiments, the EC_50_ values for viral entry into cells was calculated by fitting the luminescence data to a modified Hill plot. K_i_ values were calculated from the Hill equation. Except as noted, all plotted data points are the means of 5 or 6 independent experiments. Means ± SE were calculated.

## Supplementary Figures and Tables

**Supplementary Figure S1.**
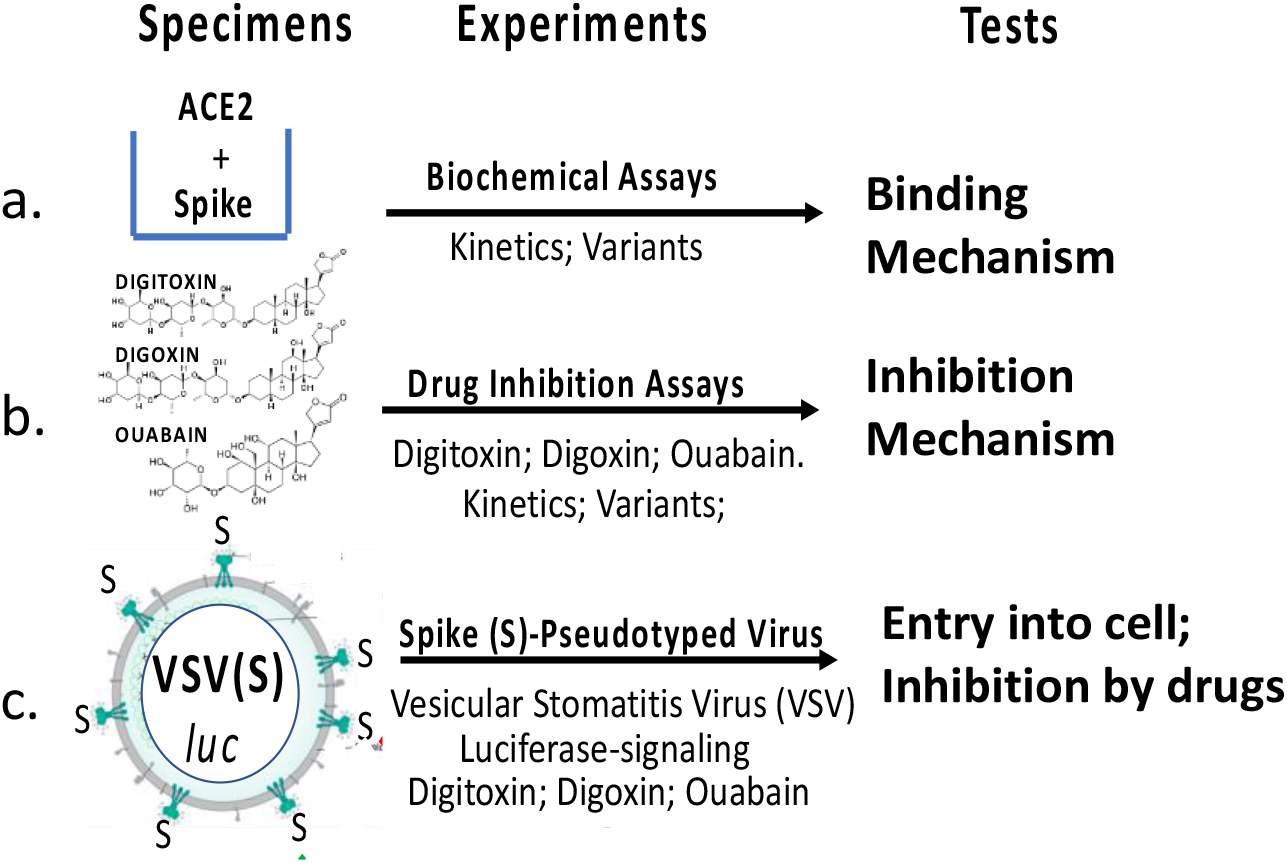
Experimental Design. **(a)** Biochemical assays. **(b)** Drug inhibition assays. **(c)** Spike pseudotyped virus assays.

**Supplementary Figure S2.**
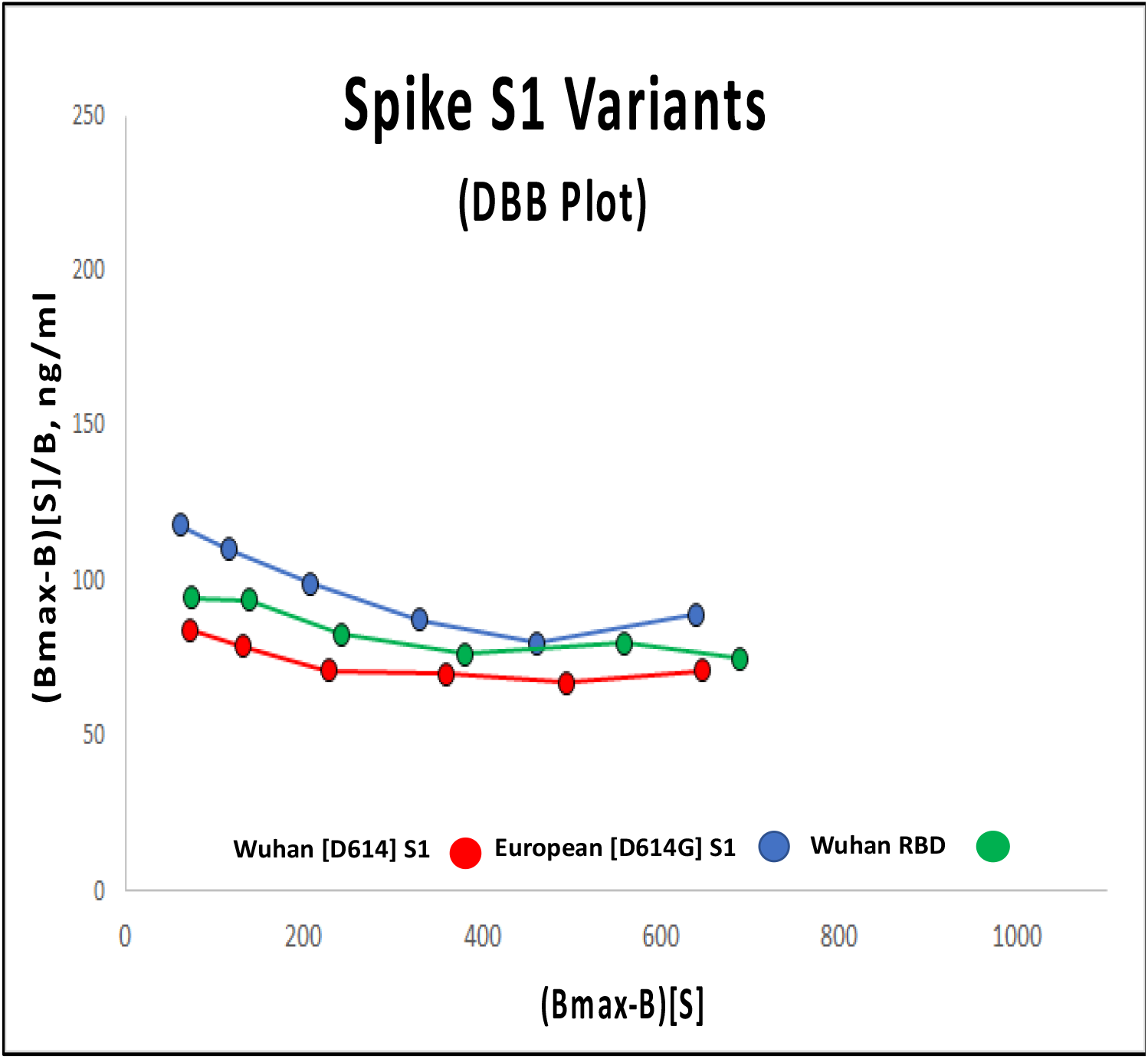
Distinguishing positive cooperativity from autocatalysis using the Dhatt, Banerjee and Battacharrya (DBB) plot for ACE2 binding data. Validated autocatalytic mechanisms, such as phase transitions, would be straight diagonal lines in the plot. Michaelis-Menten kinetics would be horizontal lines. Cooperative mechanisms would be non-linear. Color codes: Wuhan [D614] S1 (red); European [D614G] S1 (blue); Wuhan RBD (green). Each point is the average ± SE for N=5-6 independent experiments.

**Supplementary Figure S3.**
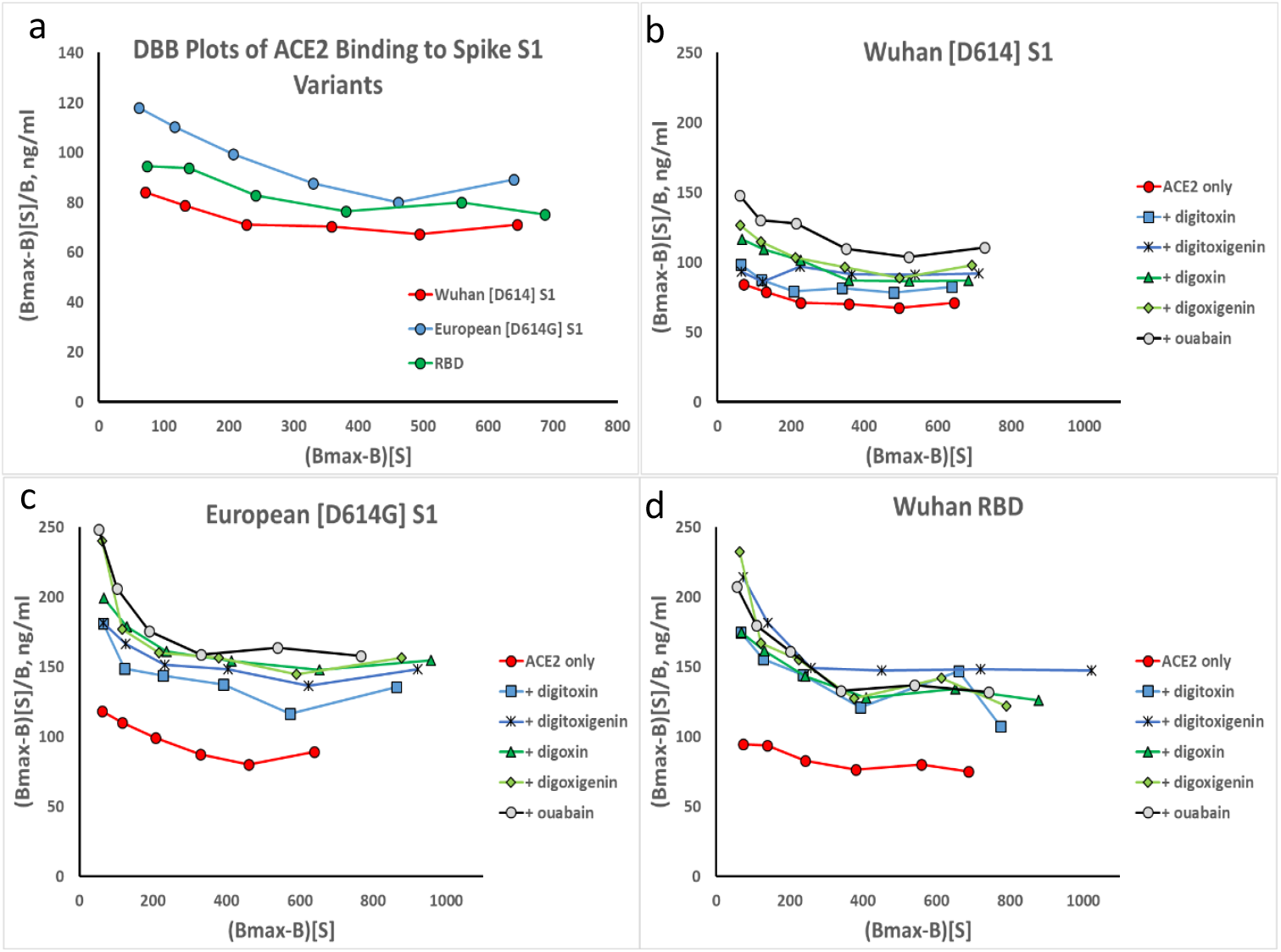
DBB plots for inhibition of ACE2 binding to spike proteins by sugar-free cardiac glycosides. **(a)** ACE2 binding to Spike variants. (**b**) ACE2 binding to Wuhan [D614] S1 inhibited by 30nM digitoxigenin and digoxigenin, and cardiac glycosides. (**c**) & (**d**): Plots for Wuhan [D614] RBD and European [D614G] S1, respectively. Red color = ACE2 alone. Consistently, elevation on the graph indicates lower K_D_. Each point is the average ± SE for N=5-6 independent experiments.

**Supplementary Figure S4.**
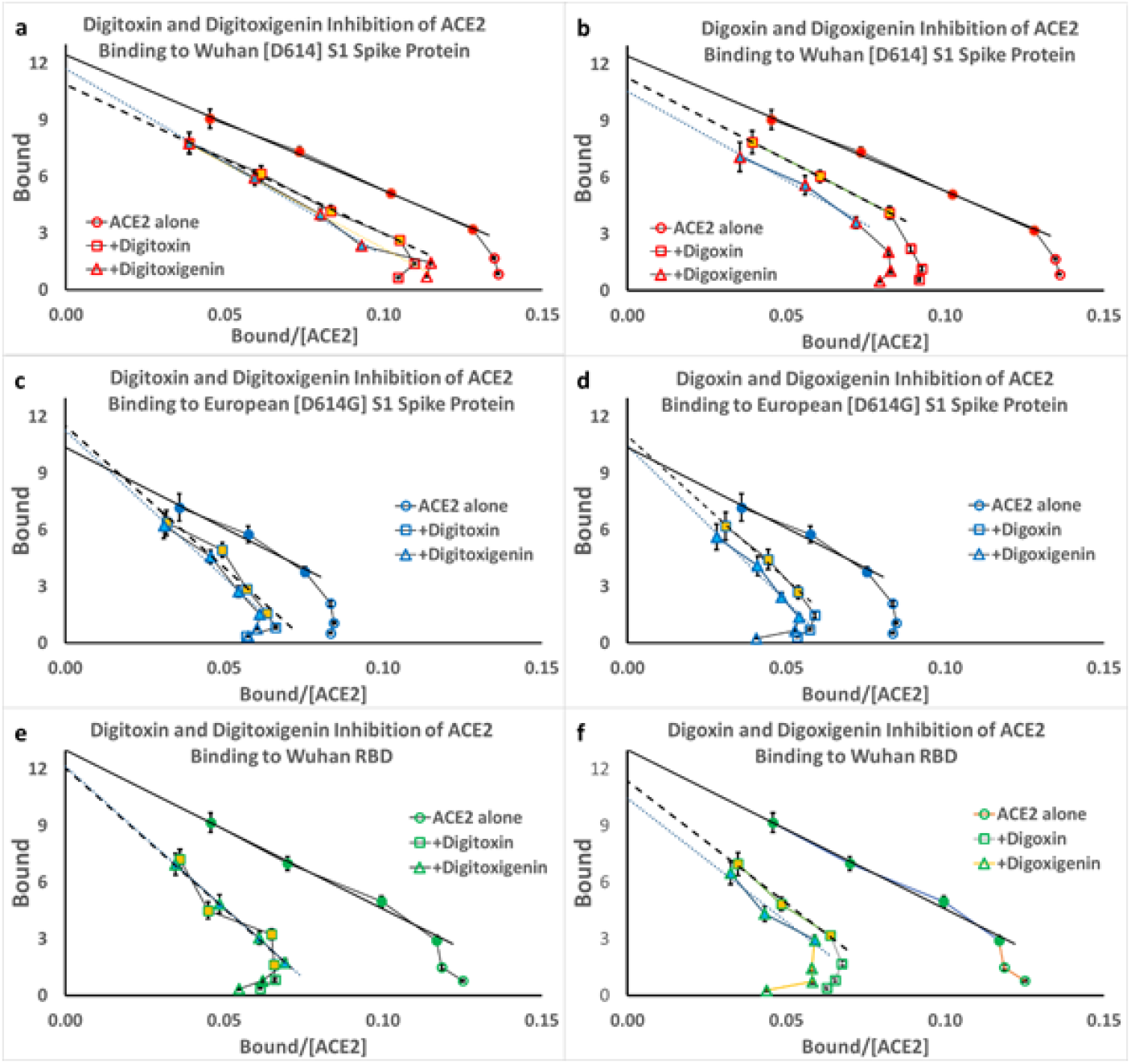
Inhibition of ACE2 binding to spike variants by digitoxigenin and digoxigenin. (**a** and **b**) Inhibition of ACE2 binding to Wuhan S1. (**c** and **d**) inhibition of ACE2 binding to European mutant [D614G] S1. (**e** and **f**) inhibition of ACE2 binding to Wuhan RBD. Each point is the average ± SE for N=5-6 independent experiments.

**Supplementary Figure S5.**
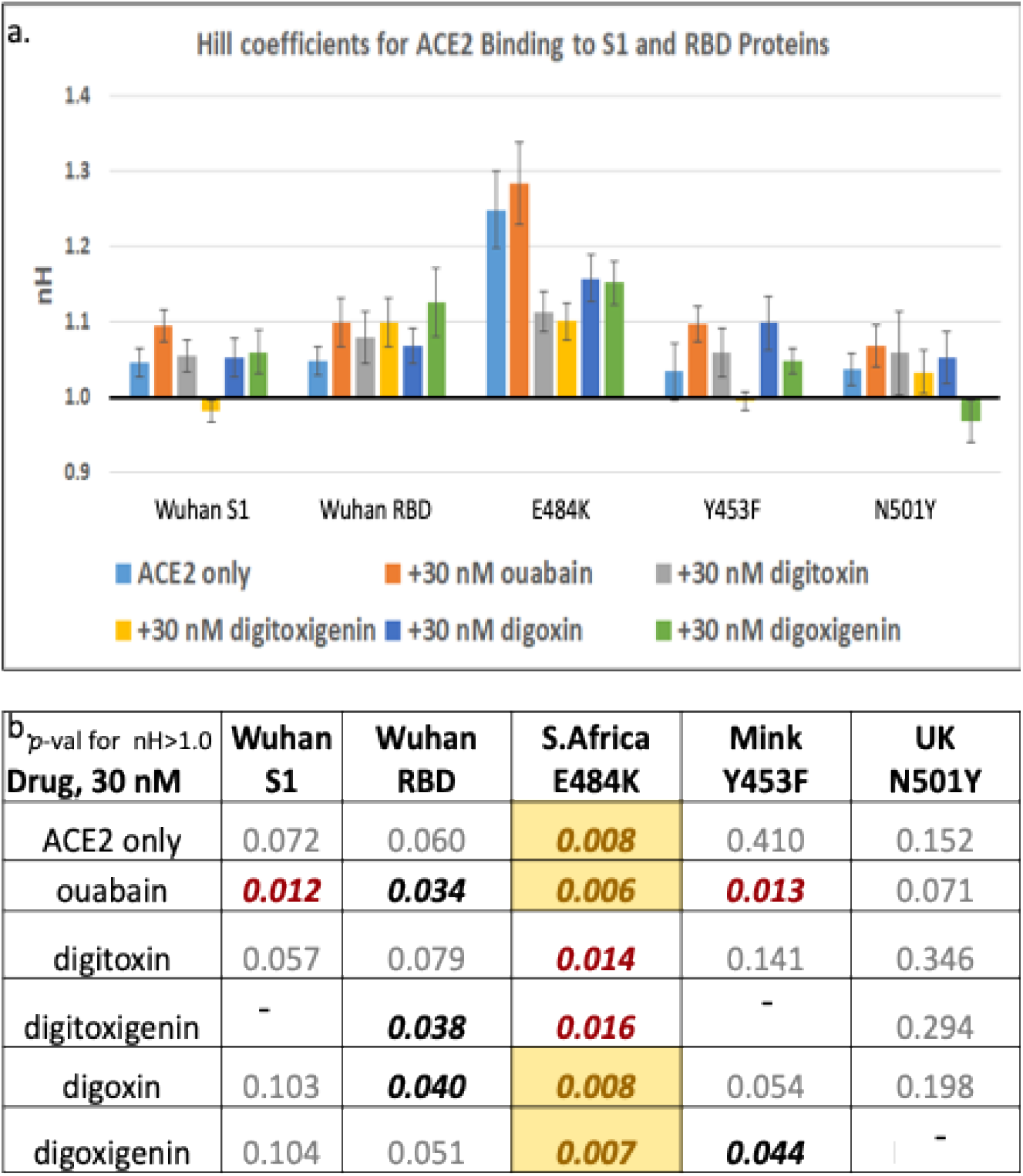
Hill coefficients for cardiac glycoside inhibition of mutant RBD proteins. (a) Values of the slope, n_H_, from Hill plots have been calculated for each of the conditions. A value of 1.0 for n_H_ means that there is no cooperativity. A value below 1.0 indicates negative cooperativity. **(b)** *p***-value for the difference of each condition from an n_H_ value of 1.0.** Color code: the color coding is **bold <0.05; bold red <0.02; yellow background <0.01**.

**Supplementary Figure S6.**
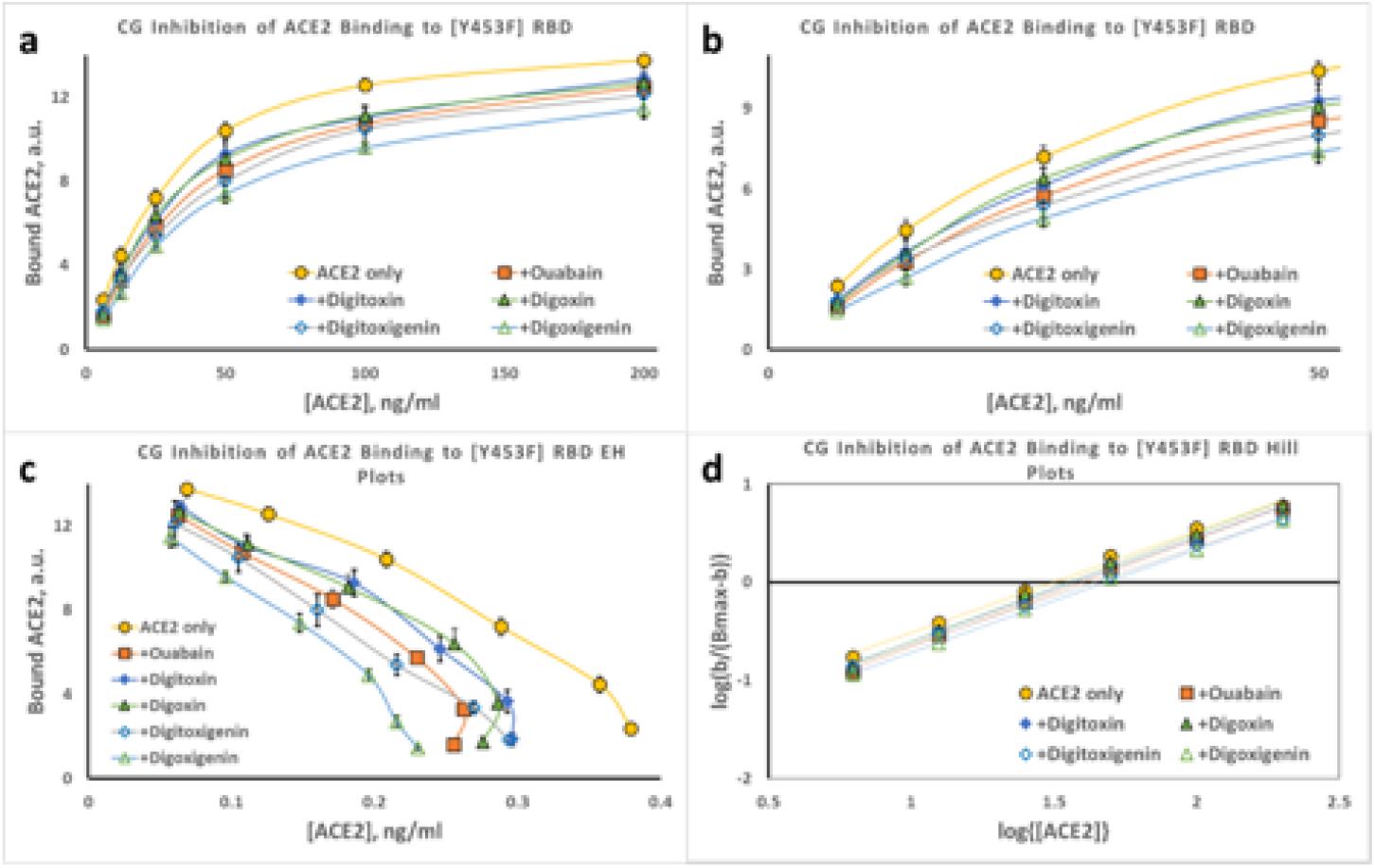
Inhibition of ACE2 binding to Mink [Y453F] RBD protein by cardiac glycoside drugs. **(a)** Substrate-Binding plots for inhibition of ACE2 binding to Mink [Y453F] RBD protein by digitoxin, digoxin, ouabain, digitoxigenin and digoxigenin (30nM). (**b**) Data from (**a**) at low concentrations of ACE2. (**c**) Eadie-Hoffstee (EH) plots of data in part (**a**). Hill plots for each titration from part (**a**). Magnitude and error for Hill coefficient (n_H_) given in **Supplemental Figure 5**. Each point is the average ± SE for N = 5-6 independent experiments.

**Supplementary Figure S7.**
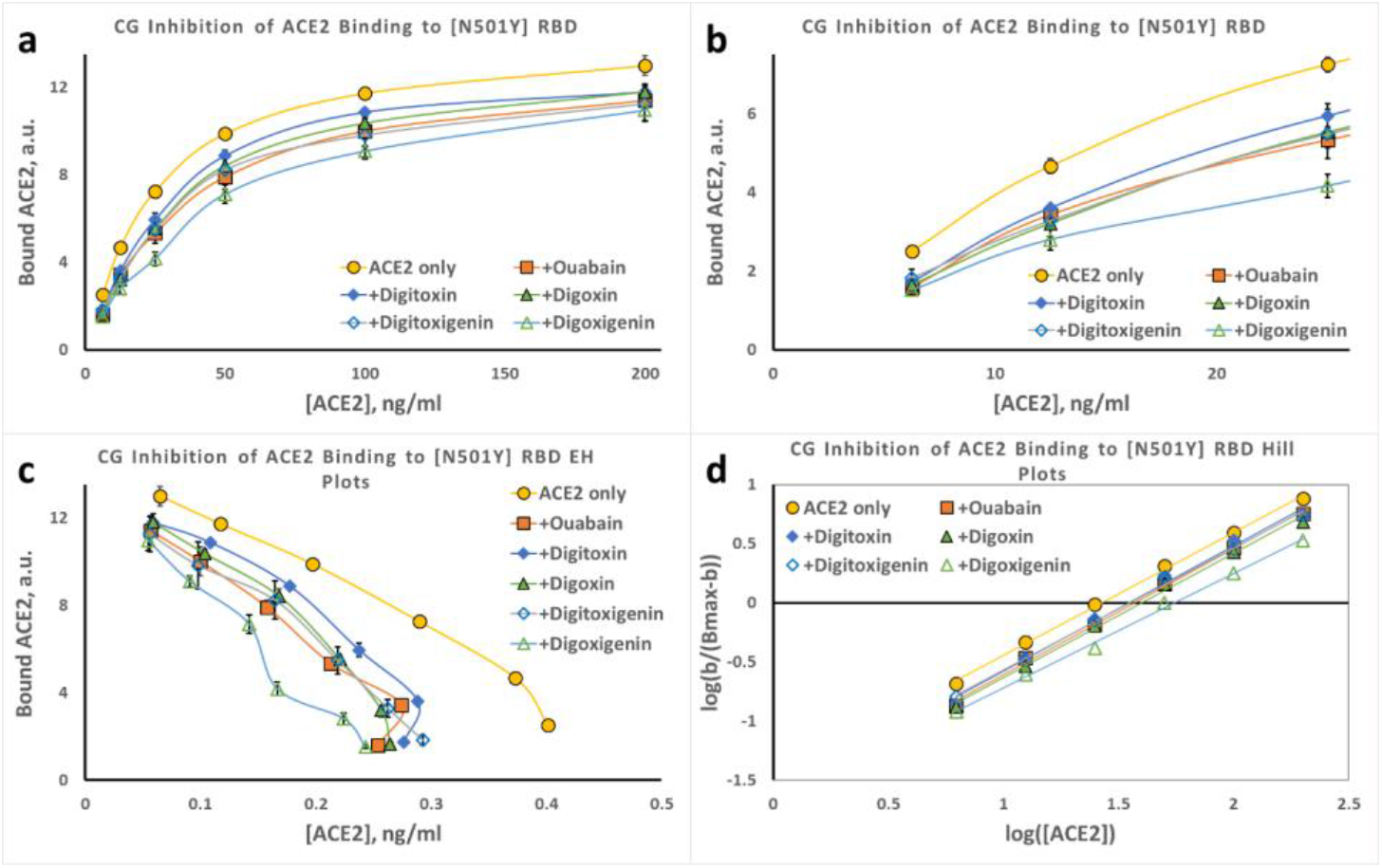
Inhibition of ACE2 binding to UK [N501]] RBD protein by cardiac glycoside drugs. **(a)** Substrate-Binding plots for inhibition of ACE2 binding to UK [N501Y] RBD protein by digitoxin, digoxin, ouabain, digitoxigenin and digoxigenin (30nM). **(b)** Data from **a** at low concentrations of ACE2. **(c)** Eadie-Hoffstee (EH) plots of data in part **a**. Hill plots for each titration from part **a**. Magnitude and error for Hill coefficient (n_H_) given in **Supplemental Figure 5**. Each point is the average ± SE for N = 5-6 independent experiments.

**Supplementary Figure S8.**
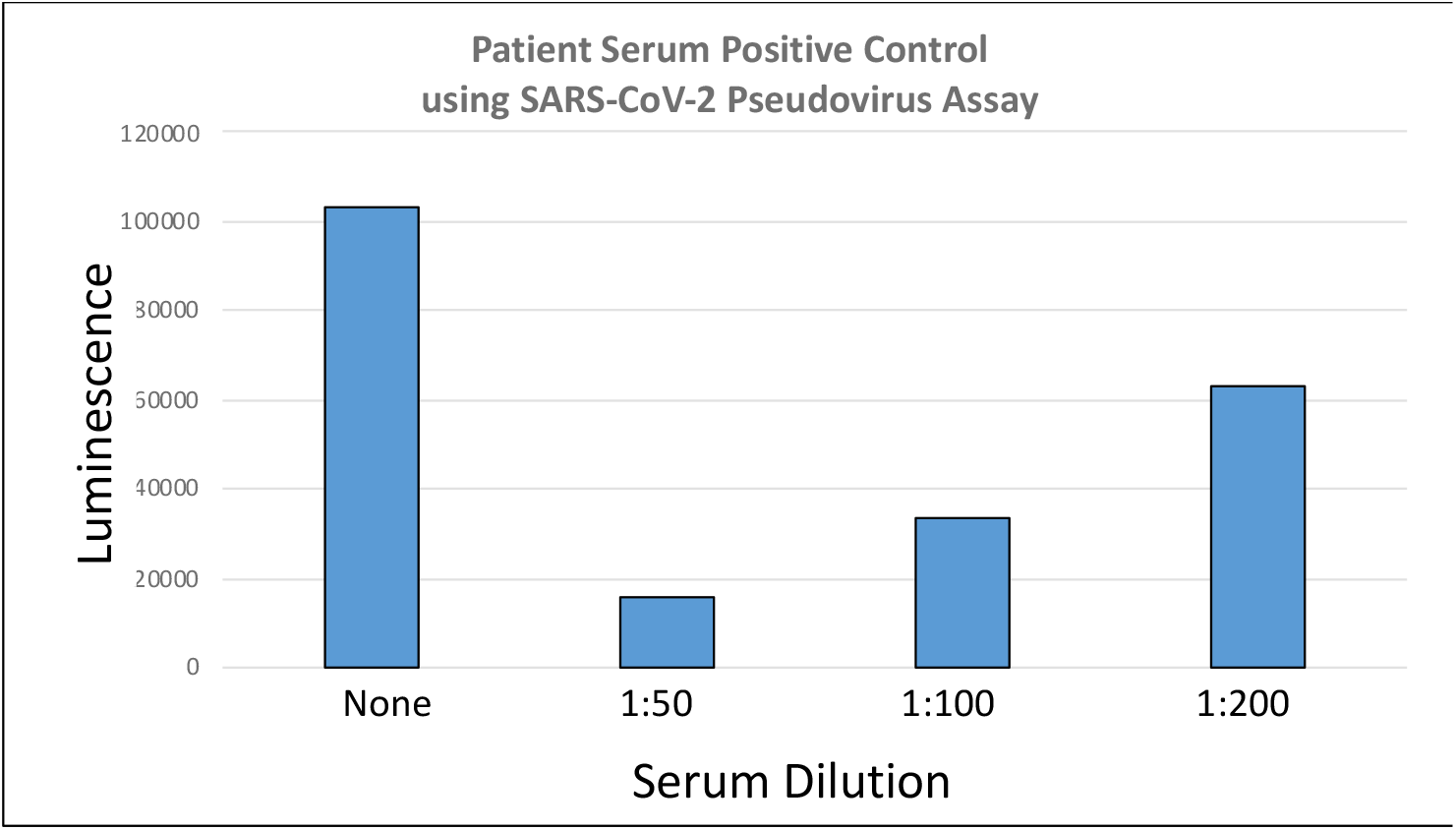
Patient-positive-control using SARS-COV-2 pseudovirus assay. Serum samples were diluted in DMEM and treated as described in methods by the triple tandem time-of-addition method. Control data are summarized in **Supplementary Table 2**.

**Supplementary Table S1.** Pseudovirus and Convalescent Serum Controls.

| Type of Control | Type of additive | Luminescence, average | Standard Deviation (N=3) |
| --- | --- | --- | --- |
| Positive | No Drugs | 105,192 | 2,352 |
| Negative | No Pseudovirus | 1205 | 195 |
| Negative | No human ACE2 | 155 | 66 |

b.
| Serum Dilution Factor | Luminescence | Luminescence minus Mock-Infection Background |
| --- | --- | --- |
| None added | 104,086 | 103,334 |
| 1:50 | 16,858 | 15,653 |
| 1:100 | 34,869 | 33,664 |
| 1:200 | 64,205 | 63,000 |

## Acknowledgements

We gratefully acknowledge support for this project by the Cooperative Health Initiative Research Program (CHIRP), supported by NHLBI/NIH (IAA-AA-HL-14-007; H.B. Pollard, PI); and by the Consortium for Health and Military Performance (CHAMP), supported by Warfighter Readiness: Optimizing Human Performance (HU00011920047; MEM-91-10314; P. Deuster, PI). We thank Laiman Tavedi for expert technical and administrative support. We thank Bette S. Pollard for her prediction that cardiac glycosides might block the interaction between ACE2 and SARS-CoV-2 spike protein, thereby preventing virus propagation, infection and COVID-19, and thank her for critically reading the manuscript.

The opinions, interpretations, conclusions and recommendations are those of the authors and are not necessarily endorsed by the U.S. Army, Department of Defense, the U.S. Government or the Uniformed Services University of the Health Sciences. The use of trade names does not constitute an official endorsement or approval of the use of such reagents or commercial hardware or software. This document may not be cited for purposes of advertisement.

## Contributions

HP, OE, TC, HC, and QY designed experiments. HC, TC, QY performed experiments. OE, AB, NW, HC and HP contributed to data analysis. HP, HC and OE wrote the paper. All co-authors critically read the paper.

## References

1 Huang, C. et al. Clinical features of patients infected with 2019 novel coronavirus in Wuhan, China. Lancet 395, 497–506, doi:10.1016/s0140-6736(20)30183-5 (2020).

2 Zhu, N. et al. A Novel Coronavirus from Patients with Pneumonia in China, 2019. The New England journal of medicine 382, 727–733, doi:10.1056/NEJMoa2001017 (2020).

3 Zhou, P. et al. A pneumonia outbreak associated with a new coronavirus of probable bat origin. Nature 579, 270–273, doi:10.1038/s41586-020-2012-7 (2020).

4 Lu, R. et al. Genomic characterisation and epidemiology of 2019 novel coronavirus: implications for virus origins and receptor binding. Lancet 395, 565–574, doi:10.1016/s0140-6736(20)30251-8 (2020).

5 Li, F., Li, W., Farzan, M. & Harrison, S. C. Structure of SARS coronavirus spike receptor-binding domain complexed with receptor. Science (New York, N.Y.) 309, 1864–1868, doi:10.1126/science.1116480 (2005).

6 Hoffmann, M. et al. SARS-CoV-2 Cell Entry Depends on ACE2 and TMPRSS2 and Is Blocked by a Clinically Proven Protease Inhibitor. Cell 181, 271–280.e278, doi:10.1016/j.cell.2020.02.052 (2020).

7 Wrapp, D. et al. Cryo-EM structure of the 2019-nCoV spike in the prefusion conformation. Science (New York, N.Y.) 367, 1260–1263, doi:10.1126/science.abb2507 (2020).

8 Walls, A. C. et al. Structure, Function, and Antigenicity of the SARS-CoV-2 Spike Glycoprotein. Cell 181, 281–292.e286, doi:10.1016/j.cell.2020.02.058 (2020).

9 Casalino, L. et al. Beyond Shielding: The Roles of Glycans in the SARS-CoV-2 Spike Protein. ACS Cent Sci 6, 1722–1734, doi:10.1021/acscentsci.0c01056 (2020).

10 Daly, J. L. et al. Neuropilin-1 is a host factor for SARS-CoV-2 infection. Science (New York, N.Y.) 370, 861–865, doi:10.1126/science.abd3072 (2020).

11 Cantuti-Castelvetri, L. et al. Neuropilin-1 facilitates SARS-CoV-2 cell entry and infectivity. Science (New York, N.Y.) 370, 856–860, doi:10.1126/science.abd2985 (2020).

12 Korber, B., Fischer B & Gnanakaran S. Tracking changes in SARS-CoV-2 Spike: evidence that D614G increases infectivity of the COVID-19 virus. Cell doi:DOI: 10.1016/j.cell.2020.06.043 (2020).

13 Becerra-Flores, M. & Cardozo, T. SARS-CoV-2 viral spike G614 mutation exhibits higher case fatality rate. Int J Clin Pract 74, e13525, doi:10.1111/ijcp.13525 (2020).

14 Zhang, J. et al. Structural impact on SARS-CoV-2 spike protein by D614G substitution. Science (New York, N.Y.), doi:10.1126/science.abf2303 (2021).

15 Zhang, L. et al. SARS-CoV-2 spike-protein D614G mutation increases virion spike density and infectivity. Nature communications 11, 6013, doi:10.1038/s41467-020-19808-4 (2020).

16 Kupferschmidt, K. Fast-spreading U.K. virus variant raises alarms. Science (New York, N.Y.) 371, 9–10, doi:10.1126/science.371.6524.9 (2021).

17 van Dorp, L. et al. Recurrent mutations in SARS-CoV-2 genomes isolated from mink point to rapid host-adaptation. bioRxiv, doi:10.1101/2020.11.16.384743 (2020).

18 Toovey, O. T. R., Harvey, K. N., Bird, P. W. & Tang, J. W. W. Introduction of Brazilian SARS-CoV-2 484K.V2 related variants into the UK. J Infect, doi:10.1016/j.jinf.2021.01.025 (2021).

19 Riva, L. et al. Discovery of SARS-CoV-2 antiviral drugs through large-scale compound repurposing. Nature 586, 113–119, doi:10.1038/s41586-020-2577-1 (2020).

20 Yuan, S. et al. Discovery of the FDA-approved drugs bexarotene, cetilistat, diiodohydroxyquinoline, and abiraterone as potential COVID-19 treatments with a robust two-tier screening system. Pharmacol Res 159, 104960, doi:10.1016/j.phrs.2020.104960 (2020).

21 Gordon, D. E. et al. A SARS-CoV-2 protein interaction map reveals targets for drug repurposing. Nature 583, 459–468, doi:10.1038/s41586-020-2286-9 (2020).

22 Cho, J. et al. Antiviral activity of digoxin and ouabain against SARS-CoV-2 infection and its implication for COVID-19. Scientific reports 10, 16200, doi:10.1038/s41598-020-72879-7 (2020).

23 Burkard, C. et al. ATP1A1-mediated Src signaling inhibits coronavirus entry into host cells. J Virol 89, 4434–4448, doi:10.1128/jvi.03274-14 (2015).

24 Ko, M. et al. Screening of FDA-approved drugs using a MERS-CoV clinical isolate from South Korea identifies potential therapeutic options for COVID-19. bioRxiv, doi:10.1101/2020.02.25.965582 (2020).

25 Wei, T. et al. in silico screening of potential spike glycoprotein inhibitors of SARS-CoV-2 with drug repurposing strategy. Research Square, doi:10.21203/rs.3.rs-17720/v1 (2020).

26 Aanouz, I. et al. Moroccan Medicinal plants as inhibitors against SARS-CoV-2 main protease: Computational investigations. J Biomol Struct Dyn, 1–9, doi:10.1080/07391102.2020.1758790 (2020).

27 Selden, R. & Smith, T. W. Ouabain pharmacokinetics in dog and man. Determination by radioimmunoassay. Circulation 45, 1176–1182, doi:10.1161/01.cir.45.6.1176 (1972).

28 Zeitlin, P. L. et al. Digitoxin for Airway Inflammation in Cystic Fibrosis: Preliminary Assessment of Safety, Pharmacokinetics, and Dose Finding. Annals of the American Thoracic Society 14, 220–229, doi:10.1513/AnnalsATS.201608-649OC (2017).

29 Hoffman, B. J. & Bigger, J. Digitalis and Allied Cardiac Glycosides Goodman and Gilman’s The Pharmacological Basis of Therapeutics Eighth Edition page 833 (Permagon Press, 1990).

30 Cornish-Bowden, A. Understanding allosteric and cooperative interactions in enzymes. Febs j 281, 621–632, doi:10.1111/febs.12469 (2014).

31 Dhatt, S., Banerjee, K. & Bhattacharyya, K. Can we distinguish positive cooperativity from autocatalysis in enzyme kinetics. J. Indian Chemical Soc 95, 909–916 (2018).

32 Lu, J. & Sun, P. D. High affinity binding of SARS-CoV-2 spike protein enhances ACE2 carboxypeptidase activity. The Journal of biological chemistry 295, 18579–18588, doi:10.1074/jbc.RA120.015303 (2020).

33 Shang, J. et al. Structural basis of receptor recognition by SARS-CoV-2. Nature 581, 221–224, doi:10.1038/s41586-020-2179-y (2020).

34 Koshland, D. E., Jr., Némethy, G. & Filmer, D. Comparison of experimental binding data and theoretical models in proteins containing subunits. Biochemistry 5, 365–385, doi:10.1021/bi00865a047 (1966).

35 Whitehead, E. P. Cooperativity and the methods of plotting binding and steady-state kinetic data. Biochem J 171, 501–504, doi:10.1042/bj1710501 (1978).

36 Monod, J., Wyman, J. & Changeux, J. P. On the nature of allosteric transitions: A plausible model J Mol Biol 12, 88–118, doi:10.1016/s0022-2836(65)80285-6 (1965).

37 Whittington, A. C. et al. Dual allosteric activation mechanisms in monomeric human glucokinase. Proceedings of the National Academy of Sciences of the United States of America 112, 11553–11558, doi:10.1073/pnas.1506664112 (2015).

38 Gohara, D. W. & Di Cera, E. Allostery in trypsin-like proteases suggests new therapeutic strategies. Trends in biotechnology 29, 577–585, doi:10.1016/j.tibtech.2011.06.001 (2011).

39 Shang, J. et al. Cell entry mechanisms of SARS-CoV-2. Proceedings of the National Academy of Sciences of the United States of America 117, 11727–11734, doi:10.1073/pnas.2003138117 (2020).

40 Yan, R. et al. Structural basis for the recognition of SARS-CoV-2 by full-length human ACE2. Science (New York, N.Y.) 367, 1444–1448, doi:10.1126/science.abb2762 (2020).

41 Sharma, S. et al. ACE2 homo-dimerization, human genomic variants and interaction of host proteins explains high population specific differences in outcomes of COVID-19. bioRxiv, doi:10.1101/2020.04.24.050534 (2020).

42 Amarelle, L. & Lecuona, E. The Antiviral Effects of Na,K-ATPase Inhibition: A Minireview. International journal of molecular sciences 19, doi:10.3390/ijms19082154 (2018).

43 Pollard, B. S., Blanco, J. C. & Pollard, J. R. Classical Drug Digitoxin Inhibits Influenza Cytokine Storm, With Implications for Covid-19 Therapy. In vivo (Athens, Greece) 34, 3723–3730, doi:10.21873/invivo.12221 (2020).

44 Yang, Q. et al. Gene therapy-emulating small molecule treatments in cystic fibrosis airway epithelial cells and patients. Respiratory research 20, 290, doi:10.1186/s12931-019-1214-8 (2019).

45 Pollard, B. S. et al. Digitoxin Inhibits Epithelial-to-Mesenchymal-Transition in Hereditary Castration Resistant Prostate Cancer. Front Oncol 9, 630, doi:10.3389/fonc.2019.00630 (2019).

46 Miller, S. C. et al. Identification of known drugs that act as inhibitors of NF-kappaB signaling and their mechanism of action. Biochemical pharmacology 79, 1272–1280, doi:10.1016/j.bcp.2009.12.021 (2010).

47 Yang, Q. et al. Cardiac glycosides inhibit TNF-alpha/NF-kappaB signaling by blocking recruitment of TNF receptor-associated death domain to the TNF receptor. Proceedings of the National Academy of Sciences of the United States of America 102, 9631–9636, doi:10.1073/pnas.0504097102 (2005).

48 Wang, H. et al. CFTR Controls the Activity of NF-kappaB by Enhancing the Degradation of TRADD. Cellular physiology and biochemistry 40, 1063–1078, doi:10.1159/000453162 (2016).

49 Srivastava, M. et al. Digitoxin mimics gene therapy with CFTR and suppresses hypersecretion of IL-8 from cystic fibrosis lung epithelial cells. Proceedings of the National Academy of Sciences of the United States of America 101, 7693–7698, doi:10.1073/pnas.0402030101 (2004).

50 Mason, D. T. & Braunwald, E. Studies on digitalis. X. Effects of ouabain on forearm vascular resistance and venous tone in normal subjects and in patients in heart failure. The Journal of clinical investigation 43, 532–543, doi:10.1172/jci104939 (1964).

51 Coates, A. L., Desmond, K., Asher, M. I., Hortop, J. & Beaudry, P. H. The effect of digoxin on exercise capacity and exercising cardiac function in cystic fibrosis. Chest 82, 543–547 (1982).

52 Moss, A. J. et al. Absorption of digoxin in children with cystic fibrosis. The Journal of pediatrics 86, 295–297 (1975).

53 Selzer, A., Hultgren, H. N., Ebnother, C. L., Bradley, H. W. & Stone, A. O. Efect of digoxin on the circulation in normal man. Br Heart J 21, 335–342, doi:10.1136/hrt.21.3.335 (1959).

54 Williams, M. H., Jr., Zohman, L. R. & Ratner, A. C. Hemodynamic effects of cardiac glycosides on normal human subjects during rest and exercise. J Appl Physiol 13, 417–421, doi:10.1152/jappl.1958.13.3.417 (1958).

55 Platz, E. A. et al. A novel two-stage, transdisciplinary study identifies digoxin as a possible drug for prostate cancer treatment. Cancer discovery 1, 68–77, doi:10.1158/2159-8274.Cd-10-0020 (2011).

56 Whitt, M. A. Generation of VSV pseudotypes using recombinant ∆G-VSV for studies on virus entry, identification of entry inhibitors, and immune responses to vaccines. J Virol Methods 169, 365–374, doi:10.1016/j.jviromet.2010.08.006 (2010).

57 Nie, J. et al. Establishment and validation of a pseudovirus neutralization assay for SARS-CoV-2. Emerg Microbes Infect 9, 680–686, doi:10.1080/22221751.2020.1743767 (2020).

58 Xiong, H. L. et al. Robust neutralization assay based on SARS-CoV-2 S-protein-bearing vesicular stomatitis virus (VSV) pseudovirus and ACE2-overexpressing BHK21 cells. Emerg Microbes Infect 9, 2105–2113, doi:10.1080/22221751.2020.1815589 (2020).

59 Almahboub, S. A.et al Evaluation of Neutralizing Antibodies Against Highly Pathogenic Coronaviruses: A Detailed Protocol for a Rapid Evaluation of Neutralizing Antibodies Using Vesicular Stomatitis Virus Pseudovirus-Based Assay. Front Microbiol 11, 2020, doi:10.3389/fmicb.2020.02020 (2020).

